# Spatial Atlas of the Mouse Central Nervous System at Molecular Resolution

**DOI:** 10.1101/2022.06.20.496914

**Authors:** Hailing Shi, Yichun He, Yiming Zhou, Jiahao Huang, Brandon Wang, Zefang Tang, Peng Tan, Morgan Wu, Zuwan Lin, Jingyi Ren, Yaman Thapa, Xin Tang, Albert Liu, Jia Liu, Xiao Wang

**Author notes:** These authors contributed equally to this work: Hailing Shi, Yichun He, and Yiming Zhou.

## Abstract

Spatially charting molecular cell types at single-cell resolution across the three-dimensional (3D) volume of the brain is critical for illustrating the molecular basis of the brain anatomy and functions. Single-cell RNA sequencing (scRNA-seq) has profiled molecular cell types in the mouse brain^1, 2^, but cannot capture their spatial organization. Here, we employed an *in situ* sequencing technique, STARmap PLUS^3, 4^, to map more than one million high-quality cells across the whole adult mouse brain and the spinal cord, profiling 1,022 genes at subcellular resolution with a voxel size of 194 X 194 X 345 nm in 3D. We developed computational pipelines to segment, cluster, and annotate 231 molecularly defined cell types and 64 tissue regions with single-cell resolution. To create a transcriptome-wide spatial atlas, we further integrated the STARmap PLUS measurements with a published scRNA-seq atlas^1^, imputing 11,844 genes at the single-cell level. Finally, we engineered a highly expressed RNA barcoding system to delineate the tropism of a brain-wide transgene delivery tool, AAV-PHP.eB^5, 6^, revealing its single-cell resolved transduction efficiency across the molecular cell types and tissue regions of the whole mouse brain. Together, our datasets and annotations provide a comprehensive, high-resolution single-cell resource that integrates a spatial molecular atlas, cell taxonomy, brain anatomy, and genetic manipulation accessibility of the mammalian central nervous system (CNS).

## Introduction

Deciphering spatial arrangements of molecular cell types at single-cell resolution in the nervous system is fundamental for understanding the molecular architecture of its anatomy, function, and disorders. scRNA-seq has revealed the complexity and diversity of cell-type composition in the mouse brain^1, 2^, yet retains little or no spatial information. Emerging spatial transcriptomic methods have shed light on the molecular organization of mouse brains^7–9^. However, existing datasets either have limited spatial resolution (100 µm)^10^—hindering *bona fide* single-cell analysis—or are restricted to particular brain subregions^11–13^. A single-cell resolved, large-scale spatial atlas across the entire CNS is highly desirable to unveil molecularly defined cell-type architectures.

Besides characterizing the spatial arrangement of molecular cell types, assessing the accessibility of genetic tools to different cell types across the brain is important to neuroscience studies and gene therapies. Specifically, engineered recombinant adeno-associated virus (rAAV) is one of the most widely used viral carriers to deliver genetic materials to targeted cell types. One of its variants, PHP.eB, can efficiently cross the blood-brain barrier (BBB) after systemic administration, enabling brain-wide transgene expression and gene therapy development^5, 6, 14^. However, the cell-type and tissue-region tropisms of AAV-PHP.eB have not been resolved nor registered in the context of the spatial brain atlas, due to the lack of robust and scalable viral characterization strategies among all the cell types and tissue regions^15^.

Here, we developed a highly efficient RNA barcoding system and combined it with a subcellular resolution, spatially resolved *in situ* RNA sequencing technique, STARmap PLUS^3, 4^, to detect 1,022 endogenous genes and PHP.eB RNA barcodes in 20 transduced CNS tissue slices at a voxel size of 194 X 194 X 345 nm. Using the spatial clustering computational pipeline ClusterMap^16^ and integrating with a published scRNA-seq atlas^1^, we generated molecularly defined cell-type and tissue region maps, and imputed transcriptome-wide spatially resolved single-cell expression profiles. Finally, we charted the AAV-PHP.eB transduction landscapes across CNS-wide cell types and tissue regions with single-cell resolution (Fig. 1a). Altogether, this work presents a comprehensive mouse CNS spatial cell atlas composed of over one million cells with their transcriptome-wide gene expression profiles, spatial coordinates, and AAV-PHP.eB tropism. This work provides experimental and computational frameworks to establish a molecular spatial atlas across scales ranging from individual RNA molecules to single cells and tissue regions.

**Figure 1.**
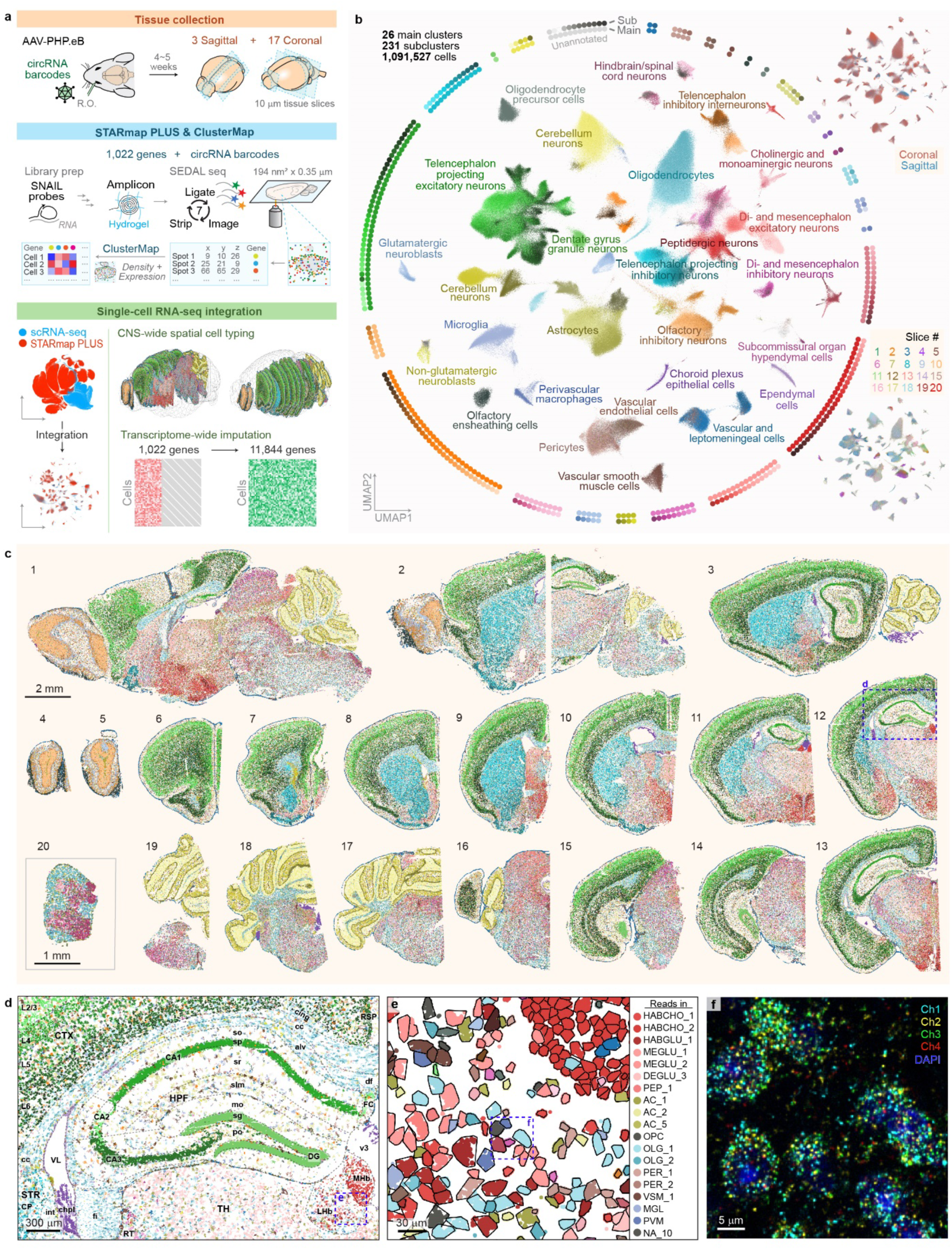
Spatial chart of molecular cell types across the adult mouse central nervous system (CNS) at subcellular resolution. **a**, Overview of the study. After systemic administration of the AAV expressing circular RNA barcodes, sagittal and coronal mouse brain tissue slices were collected. STARmap PLUS^3, 4^ was performed to detect single RNA molecules of a targeted list of 1,022 endogenous genes and trans-expressed AAV barcodes. The mRNA spot matrix was converted to a cell-by-gene expression matrix via ClusterMap^16^. By integrating existing mouse brain single-cell RNA-seq data^1^, we generated a spatial atlas across the mouse CNS. R.O., retro- orbital injection. **b**, Uniform Manifold Approximation and Projection (UMAP)^50^ of 1,091,527 cells colored by subclusters. The surrounding diagrams show 231 subclusters from 26 main clusters of cell types. Top right, the same UMAP colored by sample slice identities in coronal or sagittal directions; bottom right, UMAP colored by sample slice identity as in **c**. **c**, Spatial cell-type maps of 20 mouse CNS slices colored by subclusters. Each dot represents one cell. **d**, A zoomed-in view of tissue slice 12 in **c**. Each dot represents an mRNA amplicon color-coded by its cell-type identity. Brain regions were labeled based on the Allen Mouse Brain Reference Atlas^24^. CTX, cerebral cortex; HPF, hippocampal formation; STR: striatum; TH, thalamus; RSP: retrosplenial cortex; L2/3: layer 2/3; L4: layer 4; L5: layer 5; L6: layer 6; FC: fasciola cinerea; DG: dentate gyrus; so: stratum oriens; sp: pyramidal layer; sr: stratum radiatum; slm: stratum lacunosum-moleculare; mo: molecular layer; sg: granule cell layer; po: polymorph layer; CP: caudoputamen; RT: reticular nucleus of the thalamus; MHb: medial habenula; LHb: lateral habenula; v3: third ventricle; VL: lateral ventricle; cing: cingulum bundle; df: dorsal fornix; cc: corpus callosum; alv: alveus; fi: fimbria; int: internal capsule. **e**, A zoomed-in view of the habenula region in **d**, with cell boundaries outlined by black lines. Symbols for cell types with more than two counts were shown on the right. Cell-type abbreviations: HABCHO, habenular cholinergic neurons; MEGLU, mesencephalon excitatory neurons; DEGLU, diencephalon excitatory neurons; PEP, peptidergic neurons; AC, astrocytes; OPC, oligodendrocyte precursor cells; OLG, oligodendrocytes; PER, pericytes; VSM, vascular smooth muscle cells; MGL, microglia; PVM, perivascular macrophages; NA, unannotated. **f**, A representative fluorescent image of the highlighted square region in **e** in the first cycle of STARmap PLUS SEDAL seq. Each dot represents an amplicon generated from an RNA molecule.

### Spatial survey of molecular cell types across the adult mouse CNS

STARmap PLUS is an image-based *in situ* RNA sequencing method^3, 4^ that utilizes paired primer and padlock probes (in together termed SNAIL probes) to convert a target RNA molecule into a DNA amplicon with a gene-unique code, which enables highly multiplexed RNA detection. The DNA amplicon is further chemically modified and embedded into a hydrogel to allow robust spatial readout of the unique code by multiple rounds of sequencing by ligation (SEDAL seq) (Fig. 1a).

To achieve CNS-wide molecular cell typing, we curated a list of 1,022 genes (Supplementary Table 1, Extended Data Fig. 1a) by compiling reported cell-type marker genes from adult mouse CNS scRNA-seq datasets^1, 2, 17^. We focused on the datasets with minimal post-dissection cell-type selection to ensure that the marker genes cover all neuronal and glial cell types. A five-nucleotide code on the SNAIL probes encodes each of the 1,022 genes that can be read out by six rounds of SEDAL seq (Supplementary Table 2, Extended Data Fig. 1b). To allow orthogonal detection of AAV transcripts, we selected RNA barcode sequences from a published 25-nucleotide (nt) sequence pool^18^ without homology to the mouse transcriptome. In particular, we engineered the RNA barcode into the circular form using the autocatalytic Tornado scaffold^19^ (Extended Data Fig. 1c) to improve RNA barcode stability, allowing for extended accumulation of barcode copies in cells. Finally, one additional round of SEDAL seq was used to decode the circular RNA barcode identity (Supplementary Table 2, Extended Data Fig. 1d).

We collected STARmap PLUS datasets from twenty 10-μm-thick CNS tissue slices harvested from three mice at voxel size of 194 nm X 194 nm X 345 nm. These twenty samples included sixteen coronal brain slices, three sagittal brain slices, and one transverse slice from spinal cord lumbar segments (Supplementary Table 3, representative raw fluorescent images in Extended Data Fig. 1e). To analyze the STARmap PLUS datasets, we optimized the ClusterMap^16^ data processing workflow by adding quality control, noise reduction, and error rejection, and then generated a cell-by-gene expression matrix with RNA and cell spatial coordinates (see Methods, Extended Data Fig. 2a). In total, the datasets include over 250 million RNA reads and 1.1 million cells (Extended Data Fig. 2b).

After batch correction via Combat^20^, we pooled cells from all the tissue slices and performed cell typing by clustering single-cell RNA expression profiles hierarchically (Extended Data Fig. 2c). To annotate cell types and align with published cell-type nomenclature^1, 2, 17^, we integrated our data with an existing scRNA-seq atlas^1^ via Harmony^21^. Leiden clustering^22^ and nearest neighbor label transfer of the integrated datasets identified 26 main cell types, including 13 neuronal, 7 glial, 2 immune, and 4 vascular cell clusters (see Methods, Fig.1b). We confirmed that these main clusters exhibit corresponding canonical marker genes and expected distribution profiles across the 20 tissue slices (Extended Data Fig. 2d-e), indicating a robust annotation result. Further clustering and manual annotation within each main cluster resulted in 231 subclusters represented by 1,091,527 cells, including 190 neuronal, 2 neural crest-like glial, 14 CNS glial, 4 immune, and 9 vascular cell types (Fig. 1b, Extended Data Fig. 3, 4a-b). There are 12 unannotated subclusters (1.84% of total cells) due to the lack of marker genes for annotation (Extended Data Fig. 3l, Fig. 1b), which may result from the differences of sampling coverage between the scRNA-seq and STARmap PLUS datasets. Next, we annotated each subcluster with symbol names, cell numbers, marker genes, and spatial distributions in the anatomically defined brain regions (Supplementary Table 4). Notably, the subcluster size in our data spans approximately three orders of magnitude, ranging from abundant cell types with tens of thousands of cells such as oligodendrocytes OLG_1 (67,974 cells, 6.2% of total cells, Extended Data Fig. 3b) to rare cell types with hundreds of cells such as histaminergic neurons^23^ *Hdc*^+^ HA_1 in the posterior hypothalamus (111 cells, 0.01% of total cells, Extended Data Fig. 3i, 4c). It is worth mentioning that the cell-typing results in this study were based on the consensus between the STARmap PLUS and the published scRNA-seq datasets followed by manual annotation of participating biological experts. The STARmap PLUS dataset mapped more cells than the previous scRNA-seq dataset, potentiating more detailed cell typing and annotations in the future.

Then, we plotted molecularly defined, single-cell resolved cell-type maps throughout the adult mouse CNS (Fig. 1c). Notably, the cell-type map across the 20 CNS slices shows the clear structures for cerebral cortex (41 region-specific telencephalon projecting excitatory neuron types, TEGLU; 34 telencephalon inhibitory interneuron types, TEINH), olfactory bulb (7 olfactory inhibitory neuron types, OBINH), striatum (14 telencephalon projecting inhibitory neuron types, MSN), cerebellum (5 cerebellum neuron types; the astrocyte type AC_4), and brain stem (28 peptidergic neurons; 16 cholinergic and monoaminergic neurons; 16 di- and mesencephalon excitatory neuron types, DE/MEGLU; 8 di- and mesencephalon inhibitory neuron types, DE/MEINH; 10 hindbrain and spinal cord neuron types), fully recapitulating the anatomically defined tissue regions in the adult mouse CNS^24^ (Fig. 1c). Importantly, this molecularly defined, single-cell resolution cell-type map also reveals cell-type-specific patterns in fine tissue regions such as the medial and lateral habenula (MHb, LHb), alveus (alv), and fimbria (fi), and ependyma (Fig. 1d). A zoom-in view further illustrates diversities in cell-body sizes and cell densities across cell types (Fig. 1e). Furthermore, we envision that this molecular resolution mapping of cell transcripts and nuclear staining (Fig. 1f) could enable additional multimodal data analysis such as joint cell typing combining cell morphology and spatial transcriptomics^25^.

Remarkably, compared with previous scRNA-seq results^1, 2^, the molecular resolution, single-cell mapping across a large number of cells enables us to more precisely annotate molecular cell types by their spatial distributions. For example, among TEINH, we found a striatum-specifically located neuronal subtype, TEINH_25 (*Pvalb*^+^*Igfbp4*^+^*Gpr83*^+^) (Extended Data Fig. 4b,d). In addition, two *Th*^+^*Vip*^+^ interneuron subtypes, TEINH_10 (*Htr3a*^+^) and TEINH_22 (*Pde1c*^+^), are restrictively located in the outer plexiform layer of the olfactory bulb (OBopl) (Extended Data Fig. 4b,e), distinct from the previously identified olfactory *Th*^+^*Vip*^-^ interneurons that only reside in the glomerular layer^26^ (OBINH_7, Extended Data Fig. 4e). In addition, in brain stem excitatory neurons, besides the *Htr5b*^+^ neurons in the inferior olivary complex of the hindbrain (HBGLU_2, *C1ql1*^+^, 202 cells) reported by the scRNA-seq atlas^1^, we also identified an *Htr5b*^+^ cluster located in the habenula (HABGLU_1, *C1ql1*^-^, 318 cells) (Extended Data Fig. 3h, 4f). Interestingly, there exists a small population of TEGLU (TEGLU_36, 1,079 cells, 0.58% of TEGLU) (Extended Data Fig. 4a) with elevated expression of neuronal activity-dependent genes (*Fos* and *Junb*), while no specific behavior training or stimuli were applied to the mice prior to the brain tissue harvesting. Notably, TEGLU_36 is enriched in L2/3, which agrees with previously reported spatial patterns of its marker genes in the Allen Mouse Brain *In Situ* Hybridization (ISH) database^27^ (Extended Data Fig. 4g). This result may indicate a basal level expression of activity genes in the cortex, especially in L2/3 where intratelencephalic (IT) neurons dominate. Certainly, we also identified new spatial patterns for known glial types and new glial cell types. For example, oligodendrocyte subtypes *Opalin*^+^ OLG_1 and *Klk6*^+^ OLG_2 show preferential distribution towards the forebrain and hindbrain, respectively (Extended Data Fig. 4h). We also found that ependymal cells, instead of showing a uniform population^1^, contain two distinct subclusters both expressing the reported ependymal marker *Ccdc153*^1^ but different in the expression levels of *Fam183b* and *Hdc* (Extended Data Fig. 3d, 4i). In brief, our molecular resolution, brain-wide, large-scale *in situ* sequencing data substantially improves the spatial annotation of cell types.

### Molecularly defined tissue regions in the mouse CNS

Next, we used the brain-wide, large-scale *in situ* sequencing data to build a molecularly defined tissue region map. Such data-driven identification of tissue regions provides systematic and unbiased molecular definitions of brain tissue domains^10^. Specifically, we identified 64 tissue regions by clustering cells based on their <u>n</u>eighboring <u>c</u>ell-type <u>c</u>ompositions (NCC) using the ClusterMap pipeline^16^ (see Methods, Fig. 2a,b, Supplementary Table 5). To correlate and annotate the molecularly defined tissue regions with anatomically defined tissue regions, we registered the measured coronal slices into the previously established Allen Mouse Brain common coordinate framework (CCF, v3)^28–30^, which generated a 3D rendering of the molecularly defined tissue regions (Fig. 2c) annotated with anatomical labels (Fig. 2d, Extended Data Fig. 5).

**Figure 2.**
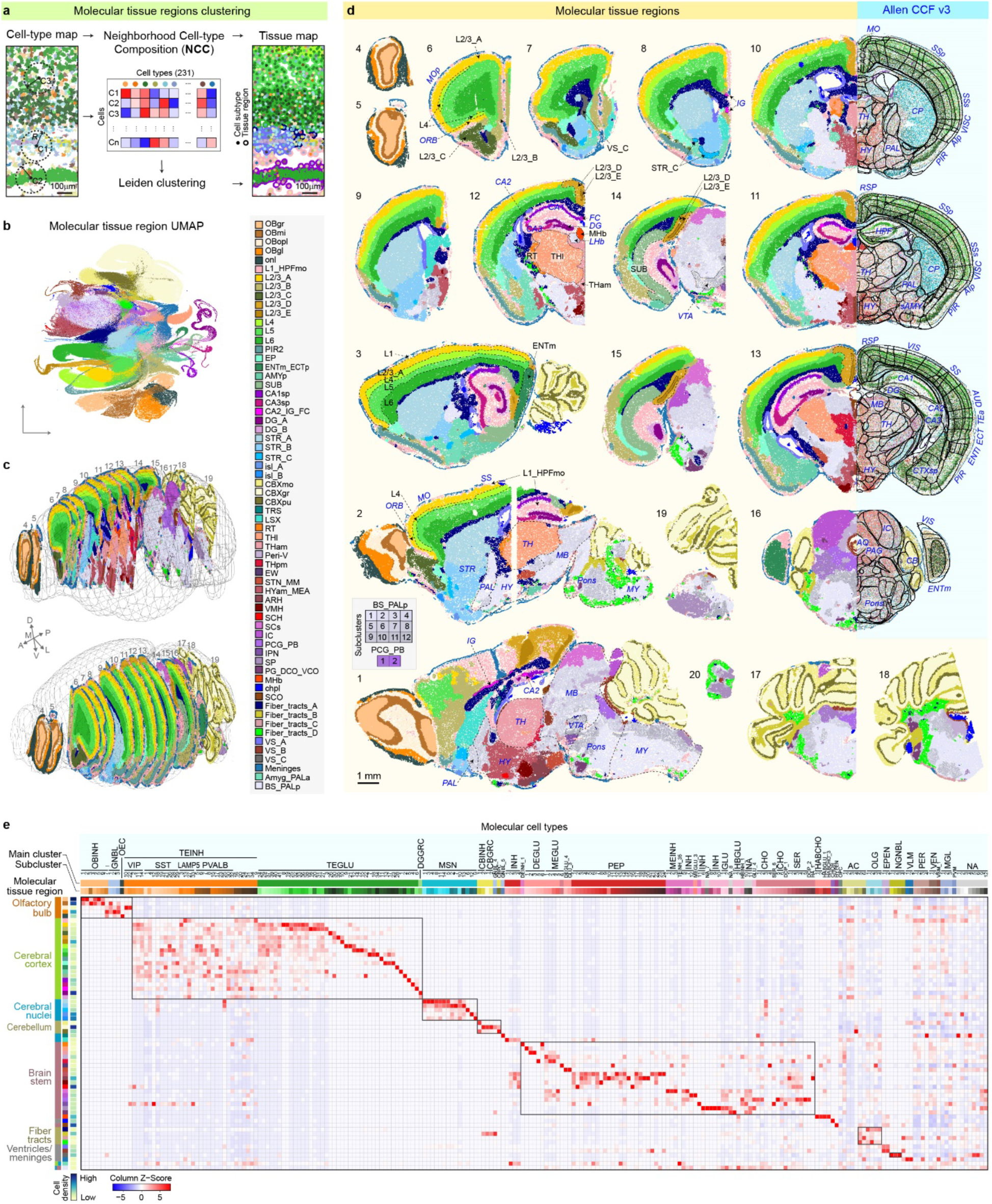
Molecular tissue regions across the adult mouse CNS. **a**, Workflow of molecular tissue region clustering. Cells are clustered based on the neighboring cell-type composition (NCC) within a 500-pixel (97.2-μm) radius circle for tissue region identification. **b**, UMAP of cell NCC vectors colored by its assigned tissue region. **c**,**d**, Molecular tissue region maps registered into the Allen Mouse Brain Common Coordinate Framework^28^ (CCF, v3, 10 μm resolution) visualized in 3D (16 coronal slices combined, **c**) and 2D (individual slices, **d**). Representative registrations were shown to compare corresponding molecularly defined tissue regions (colored by molecular tissue region identities) with anatomically defined tissue regions (colored by molecular cell types) of the same slice (**d**, right). Each dot represents a cell. Anatomically defined tissue regions were labeled in italics in blue. **e**, A heatmap showing the distribution of molecular cell types (231 subtypes) across all molecular tissue regions. Heatmap colors represent the scaled cell-type percentage composition across tissue regions. Cell density for each molecular tissue region was shown as a color bar on the left. Data are provided in the accompanying Source Data file. Abbreviations of tissue regions were labeled based on the Allen Mouse Brain Reference Atlas^24^. OBgr, olfactory bulb, granule layer; OBmi, olfactory bulb, mitral layer; OBopl, olfactory bulb, outer plexiform layer; OBgl, olfactory bulb, glomerular layer; onl, olfactory nerve layer of main olfactory bulb; L1, cortical layer 1; HPFmo, hippocampal formation, molecular layer; L2/3, cortical layer 2/3; L4, cortical layer 4; L5, cortical layer 5; L6, cortical layer 6; PIR2, piriform area, pyramidal layer; EP, endopiriform nucleus; ENT, entorhinal area; ENTm, ENT medial part; ENTl, ENT lateral part; ECTp, ectorhinal area, posterior part; SUB, subiculum; AMYp, posterior amygdala; CA1sp, field CA1, pyramidal layer; CA3sp, field CA3, pyramidal layer; IG, induseum griseum; FC, fasciola cinerea; DG, dentate gyrus; STR, striatum; isl, islands of Calleja; CB, cerebellum; CBXmo, cerebellar cortex, molecular layer; CBXgr, cerebellar cortex, granular layer; CBXpu, cerebellar cortex, Purkinje layer; TRS, triagnular nucleus of septum; LSX, lateral septal complex; RT, reticular nucleus of the thalamus; TH, thalamus; THI, lateral TH; THam, anterior- medial TH; Peri-V, peri-medial ventricles; THpm: posterior medial TH; EW, Edinger-Westphal nucleus; STN, subthalamus nucleus; MM, medial mammillary nucleus; HY, hypothalamus; HYam, anterior-medial HY; MEA, medial amygdalar nucleus; ARH, arcuate hypothalamic nucleus; VMH, ventromedial hypothalamic nucleus; SCH, hypothalamic suprachiasmatic nucleus; SCs, superior colliculus, sensory related; IC, inferior colliculus; PCG, pontine central gray; PB, parabigeminal nucleus; IPN, interpeduncular nucleus; SP, spinal cord related; PG, pontine gray; DCO, dorsal cochlear nucleus; VCO, ventral cochlear nucleus; Hb, habenula; MHb, medial Hb; LHb, lateral Hb; chpl, choroid plexus; SCO, subcommissural organ; VS, ventricular system; BS, brain stem; MB, midbrain; MY, medulla; PAL, pallidum; PALa, anterior PAL; PALp, posterior PAL; VTA, ventral tegmental area; HPF, hippocampal formation; ORB, orbital area; ACA, anterior cingulate area; MO, somatomotor areas; MOp, primary MO; RSP, retrosplenial area; SS, somatosensory area; SSp, primary SS; SSs, secondary SS; VISC, visceral area; AIp, agranular insular area, posterior part; sAMY, striatum-like amygdalar nuclei; VIS, visual area; AUD, auditory area; TEa, temporal association area; CTXsp, cortical subplate; AQ, cerebral aqueduct; PAG, periaqueductal gray. Cell type abbreviations: OBINH, olfactory inhibitory neurons; GNBL, glutamatergic neuroblasts; OEC, olfactory ensheathing cells; TEINH, telencephalon inhibitory interneurons; TEGLU, telencephalon excitatory neurons; DGGRC, dentate gyrus granule neurons; MSN, medium spiny neurons; CBINH, cerebellar inhibitory neurons; CBGRC, cerebellar granule neurons; CBPC, Purkinje neurons; INH, inhibitory neurons; GLU, excitatory neurons; DEGLU, diencephalon excitatory neurons; MEGLU, mesencephalon excitatory neurons; MEINH, mesencephalon inhibitory neurons; PEP, peptidergic neurons; HBGLU, hindbrain excitatory neurons; CHO, cholinergic neurons; SER, serotonergic neurons; DOP, dopaminergic neurons; HA, histaminergic neurons; HABCHO, habenular cholinergic neurons; CHOR, choroid plexus epithelial cells; HYPEN, subcommissural organ hypendymal cells; OPC, oligodendrocyte precursor cells; AC, astrocytes; OLG, oligodendrocytes; EPEN, ependymal cells; NGNBL, non- glutamatergic neuroblasts; VLM, vascular and leptomeningeal cells; VSM, vascular smooth muscle cells; MGL, microglia; VEN, vascular endothelial cells; PVM, perivascular macrophages; NA, unannotated.

Overall, the molecularly defined tissue regions align well with anatomically defined tissue regions. For example, molecular cerebral cortical regions show the same laminar organization as anatomical layers in the cerebral cortex (L1, L2/3, L4, L5, L6)^10, 13^, and they further reveal the molecular heterogeneity in cortical regionalization along the medial-lateral and anterior-posterior axes^31^ (Fig. 2d, regions L2/3_A-E across slices 1-3, 6-15, Extended Data Fig. 5b). In the hippocampal formation, molecular tissue regions correspond to CA1-3 and dentate gyrus (DG) in the hippocampal region as well as subiculum (SUB) and medial entorhinal area (ENTm) in the retrohippocampal region (Fig. 2d, slices 1-3, 11-15). Likewise, in the striatum, molecular tissue regions recapitulate known anatomical divisions, including the dorsal (region STR_A, D1 medium spiny neurons MSN_3 and MSN_4 enriched) and ventral (region STR_B, D1 MSN_2 and D2 MSN_7 enriched) areas, and islands of Calleja (regions isl_A and isl_B, D1 MSN_12 and MSN_9 enriched, respectively) (Fig. 2d, slices 2-3, 8-12, Extended Data Fig. 5c). In addition, both the molecular olfactory bulb regions (OBgr, OBmi, OBopl, OBgl) and the molecular cerebellar cortical regions (CBXmo, CBXpu, CBXgr) form delicate layered structures corresponding to the anatomically defined layers (Fig. 2d, OB: slices 1-2, 4-6, CBX: slices 1-3, 16-19; Extended Data Fig. 5d,e). Finally, multiple subdivisions of the molecular regions in the hypothalamus (HY) appear as spatially segregated nuclei structures distributed along the anterior-posterior, dorsal-ventral, and medial-lateral axes (Fig. 2d, slices 1, 12-13), similar to the corresponding anatomically defined structures (Extended Data Fig. 5f). Indeed, the match of molecularly defined tissue regions with anatomically defined tissue regions confirms the molecular basis of the anatomical organizations in the mouse brain.

Certainly, molecularly defined tissue regions are not necessarily the same as the anatomically defined tissue regions. On one hand, molecular tissue regions further illustrate cell-type-defined patterns that lack apparent anatomical borderlines. For instance, we found a previously underrecognized putative layer 4 structure (region L4, marked by *Rorb*^+^ TEGLU_9) that locates in the motor and orbital cortices extended from the somatosensory and visual areas (Fig. 2d, slices 2, 6), supported by recent characterizations of a laminar zone in the motor cortex with L4- like synaptic connections^32^ and spatial domains defined in the Visium mouse brain atlas^10^. In the striatum, we identified a unique structure (STR_C) marked by *Foxp2*^+^ D1 MSN_6 (Extended Data Fig. 5c, Fig. 2d, slices 8-11, 2-3), consistent with the spatial distribution of the atypical D1 MSNs defined by a previous MERFISH study^12^. Interestingly, we identified a core-shell-like structure in the retrosplenial cortex (RSP), presubiculum (PRE), and postsubiculum (POST), where layer 2 (region L2/3_E, marked by *Tshz2*^+^*Dkk3*^+^ TEGLU_10) is surrounded by layers 1 and 3 (region L2/3_D, marked by *Tshz2*^+^*Cbln1*^+^ TEGLU_35 or *Tshz2*^+^*Rxfp1*^+^ TEGLU_30) (Fig. 2d, slices 12- 15). On the other hand, one molecular tissue region could contain multiple anatomically defined tissue regions, which suggests shared cell-type compositions. For example, cortical layer 1 and hippocampal molecular layers share similar cell-type composition and cell density (region L1_HPFmo, Fig. 2e, Extended Data Fig. 5a). They are both occupied primarily by TEINH telencephalon inhibitory interneuron subtypes (*e.g.*, *Adarb2*^+^ TEINH_9, *Lamp5*^+^ TEINH_13) and astrocytes (*e.g.*, AC_2, AC_3) (Fig. 2d, slices 2-3, 11-15), which may be related to the homologous developmental origins of the isocortex and allocortex^31, 33^. As another example, in the hippocampal area, the CA2 pyramidal layer exhibits a high resemblance with the indusium griseum (IG) and the fasciola cinerea (FC) in molecular expression, agreeing with previous reports^34, 35^ (region CA2_IG_FC, marked by *Neurod6*^+^*Rgs14*^+^ TEGLU_6; Fig. 2d, slices 1, 8, 11- 12).

Next, we aimed to pinpoint molecular tissue region-specific cell types and gene markers. To this end, we plotted the distribution heatmap of the 231 subcluster cell types along the 64 molecular tissue regions to examine their cell-type composition signatures (Fig. 2e, Supplementary Table 5). Indeed, most molecular tissue regions can be reliably defined by marker cell types, demonstrated by the distinct correlation pattern along the heatmap diagonal (Fig. 2e). For example, molecular habenular regions revealed clear contrast between medial (MHb) and lateral habenula (LHb): MHb is enriched for cholinergic neurons (e.g., *Nwd2*^+^ HABCHO_2) while LHb is dominated by glutamatergic neurons (e.g., *Resp18*^+^*Baiap3*^+^ MEGLU_1-2) (Fig. 1e, 2e). In addition, the rest of molecular thalamic regions show distinct subdivisions outlining anterior-medial (THam), anterior-lateral (RT, reticular nucleus of the thalamus), posterior-medial (THpm), and posterior-lateral thalamus (THl) (Fig. 2d, slices 11-12, 1-2), marked by excitatory neurons DEGLU_2-3 (*Necab1*^+^), inhibitory neurons DEINH_1 (*Pvalb*^+^*Hs3st4*^+^), excitatory neurons DEGLU_5 (*Cbln4*^+^*Wnt4*^+^), and excitatory neurons DEGLU_1 (*Prkcd*^+^*Synpo2*^+^) and DEGLU_6 (*Prox1*^+^*Cbln1*^+^), respectively (Fig. 2e). Particularly, in the ventricular area (Extended Data Fig. 5g), the molecular region VS_C in the subventricular zone (SVZ) is marked by non-glutamatergic neuroblasts NGNBL_1-3 (*Mki67*^+^, *Cdca7*^+^, and *Meis2*^+^, respectively), reflecting the presence of immature neurons in SVZ^36^. Moreover, by focusing on cell-type distribution across molecular tissue regions, we observed that different cell types show diverse degrees of regional localization (Fig. 2e). For example, most glutamatergic neuronal cell types (TEGLU, DEGLU, MEGLU, HBGLU, *etc.*) show region-specific distributions (Fig. 2e), consistent with previous reports^1, 10, 31^. Specifically, 16 out of the 19 molecular cerebral cortical regions and 9 out of 18 molecular brain stem regions are marked by glutamatergic neuronal types (Fig. 2e, Supplementary Table 5). As for inhibitory neurons, while the majority of the telencephalon interneuron subtypes (TEINH) show broad distribution across multiple molecular tissue regions, there are examples of highly region-specific inhibitory neuron subtypes, such as INH_1 (*Atp2b4*^+^*Zic1*^+^) in the lateral septal complex (LSX), DEINH_1 (*Pvalb*^+^*Hs3st4*^+^) in RT, and INH_11 (*Gad1*^+^*Chrna2*^+^) in interpeduncular nucleus (IPN) (Fig. 2e). As another example, we confirmed region-specific patterns of astrocyte subtypes^1, 37^ in the telencephalon (AC_2-3, *Mfge8*^+^), non-telencephalon (AC_1, *Agt*^+^), cerebellar cortex (AC_4, *Gdf10*^+^), and fiber tracts (AC_5, *Gfap*^+^*Mbp*^+^; AC_6, *Gfap*^+^*Myoc*^+^) (Fig. 2e, Extended Data Fig. 5h,i).

Additionally, we identified marker genes in each molecular tissue region by comparing region mean expression profiles (see Methods; Extended Data Fig. 6a, Supplementary Table 5). We recovered canonical region markers that agree with the Allen ISH database^27^, such as cortical layer markers (e.g. *Cux2* and *Satb2* in L2/3_A, *Rorb* and *Rspo1* in L4, *Fezf2* and *Scube1* in L5, and *Col5a1* and *Rprm* in L6) and hippocampal pyramidal layer markers (e.g., *Fibcd1* in CA1, *Rgs14* in CA2, *Npy2r* in CA3, and *Prox1* in DG). Importantly, we were able to capture gene signatures in very fine structures of brain nuclei, such as *Cartpt* in the Edinger-Westphal nucleus (EW), *Chrna2* in the interpeduncular nucleus (IPN), and *Rgs16* in the hypothalamic suprachiasmatic nucleus (SCH), in line with the previous literature^38–40^ (Extended Data Fig. 6b-d). In conclusion, we reported a resource of molecular tissue regions across the whole mouse CNS registered with anatomical definitions and annotated with region-specific cell types and marker genes (Supplementary Table 5).

### Imputation for transcriptome-wide spatial gene expression in the mouse CNS

To establish transcriptome-wide spatial profiling of the mouse CNS, we imputed cellular transcriptomic profiles by integrating the STARmap PLUS data with the scRNA-seq atlas^1^ by a previously reported <u>m</u>utual <u>n</u>earest <u>n</u>eighbors (MNN) imputation method^41^. Specifically, using the 1,022-gene STARmap PLUS measurements and the 11,844-gene scRNA-seq atlas as inputs, we first generated 1,022 intermediate mappings by the leave-one-(gene)-out strategy (see Methods). The resulting intermediate mappings were used to define the size of nearest neighbors (Extended Data Fig. 7a, Supplementary Table 6) and compute weights between STARmap PLUS identified cells and scRNA-seq cells for the final transcriptome-scale imputation. In total, we imputed 11,844-gene expression profiles for 1.09 million cells in the STARmap PLUS datasets as the output for the transcriptome-wide spatial cell atlas of the mouse CNS (Fig. 3a, see Methods).

**Figure 3.**
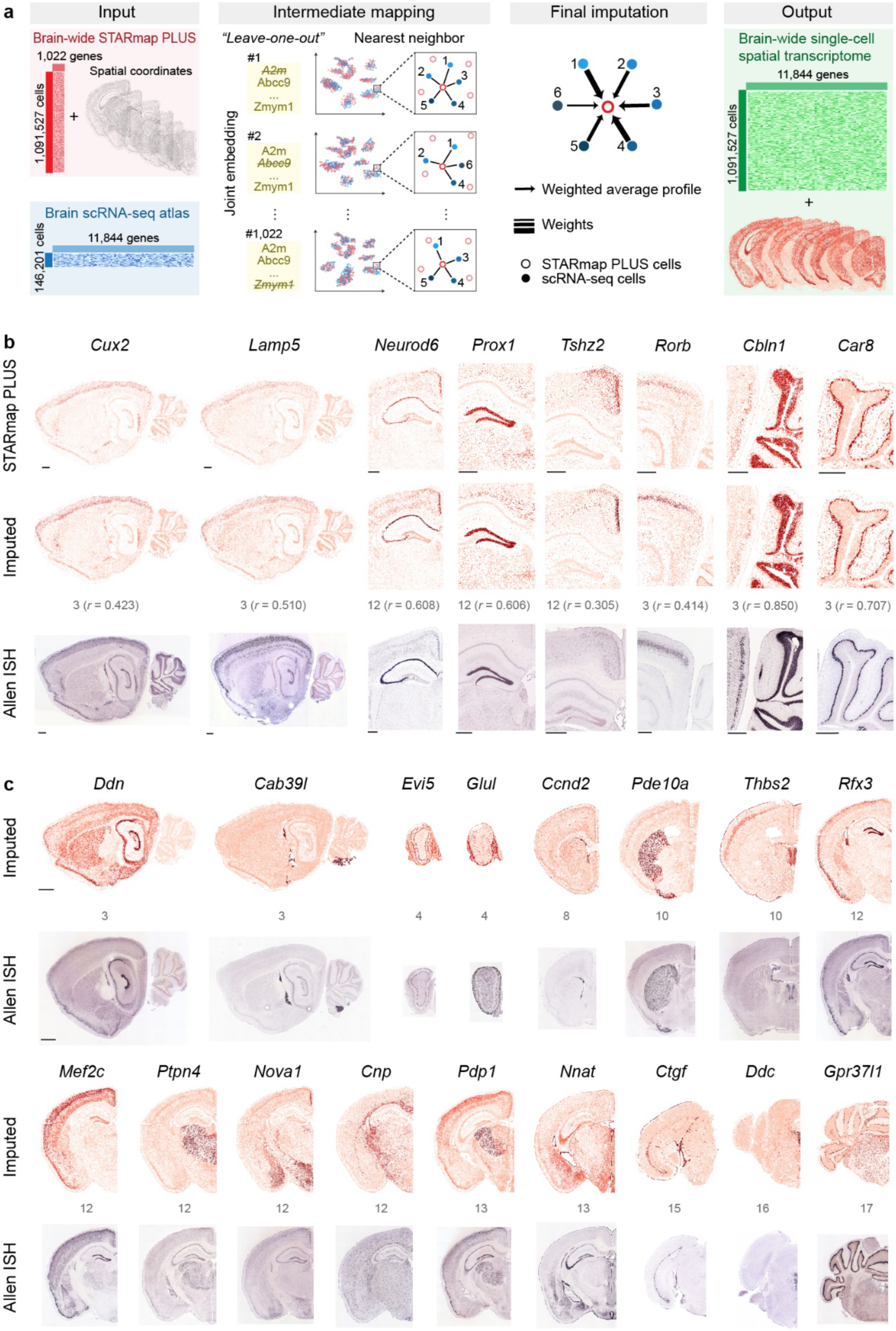
Transcriptome-scale adult mouse CNS spatial atlas by gene imputation. **a**, Schematics of the imputation workflow. Using the 1,022-gene STARmap PLUS measurements and an 11,844-gene scRNA-seq atlas^1^ as input, we first performed 1,022 intermediate mappings by the leave-one-(gene)-out strategy (see Methods). The resulting intermediate mappings were used to compute weights between STARmap PLUS identified cells and scRNA-seq cells. Finally, 11,844-gene expression profiles in STARmap PLUS identified cells were imputed as the output for the transcriptome-wide spatial cell atlas of the mouse CNS. **b**, Comparison of representative imputed spatial gene expression maps with corresponding STARmap PLUS and Allen Mouse Brain In Situ Hybridization (ISH) gene expression maps^27^. Each dot represents a cell colored by the expression level of a gene. Scale bar, 0.5 mm. The sample slice number and Pearson correlation coefficient *r* between STARmap PLUS measurements and imputed results were labeled in gray. **c**, Independent validation of imputation performance by comparing the imputed spatial expression profile of selected genes outside of the STARmap PLUS 1,022 gene list with the Allen ISH database^27^. Scale bar, 1 mm.

To validate the final imputation results, we compared the imputed results with ground-truth measurements from the STARmap PLUS and the Allen ISH database^27^. For example, regional markers show consistent spatial patterns across imputed and experimental results: *Cux2* in cortical layers 2-4, *Rorb* in cortical layer 4, *Prox1* in the dentate gyrus (DG), *Tshz2* in RSP, *Lmo3* in the piriform (PIR), *Pdyn* in the ventral striatum (STRv), *Gng4* in the olfactory bulb granular layer (OBgr), and *Hoxb6* and *Slc6a5* in the spinal cord. Moreover, cell-type markers for both abundant and rare cell types were also accurately imputed: cortical interneuron marker *Lamp5*, cerebellum neuron marker *Cbln1*, Purkinje cell maker *Car8*, and serotonergic neuron marker *Tph2* (Fig. 3b, Extended Data Fig. 7b).

After confirming the accuracy of imputed results for STARmap PLUS measured genes, we further benchmarked the imputed results of unmeasured genes with the Allen ISH database^27^. The imputed results successfully predicted the known spatial patterns of unmeasured genes (Fig. 3c).

For example, imputed spatial patterns of cell-type marker genes fully recapitulated those reported in the Allen ISH database^27^, including *Cab39l* for choroid epithelial cells (CHOR), *Cnp* for oligodendrocytes, and *Ddc* for dopaminergic neurons. Remarkably, for genes that express at different levels across multiple regions, the imputed results can quantitatively predict their relative regional expression. For instance, we correctly predicted: *Rfx3*, a transcription factor, highly expressed in DG, PIR, and choroid plexus, and modestly in cortical L2/3, DG, and ependyma; *Nova1*, an RNA-binding protein, densely in RSP L2/3, amygdala, and medial hypothalamic nuclei, and sparsely in the LHb; *Nnat*, a proteolipid, highly expressed in the ependyma, and modestly in the CA3, amygdala, and medial brain stem.

Finally, we asked whether we could uncover more tissue region-specific marker genes from the imputed results. Taking MHb as an example, in addition to the MHb-enriched genes in the 1,022- gene list (e.g., *Gm5741*, *Gng8*, *Nwd2*, and *Lrrc55*)^27, 42^, we identified 61 genes from the imputed gene list that are enriched in MHb (*Z*-score > 5, Supplementary Table 6), including *Kctd8*, *Af529169*, *Vav2*, *Lrrc3b*, *Prkcq*, and *Myo16*. Cross-validation with the Allen ISH database^27^ confirms that these genes are indeed correct MHb markers (Extended Data Fig. 7c).

Collectively, combining the molecular resolution, brain-wide, large scale STARmap PLUS datasets with the transcriptome-wide scRNA-seq atlas^1^, we generated a transcriptome-wide spatial cell atlas of the mouse CNS with single-cell resolution. We envision this imputed, expanded atlas as a valuable resource to discover spatially variable genes, spatially co-regulated gene programs, and cell-cell interactions.

### Quantitative evaluation of AAV-PHP.eB tropisms across CNS tissue regions and cell types

The rAAV has been the leading vector for *in vivo* transgene delivery and gene therapy development in neuroscience research and gene therapies, owing to its low immunogenicity, low cytotoxicity, and low risks of genome integration^43, 44^. Different AAV serotypes exhibit varied tropisms, involving differential interactions with target cell receptors through AAV capsid proteins^43, 44^. However, characterization of AAV tropism is largely limited by the inability to sample a significant population of cells with annotated cell types. A quantitative brain-wide and cell-type resolved tropism profile would be beneficial for assessing the applicability of AAV variants and providing insights into AAV capsid rational design. To this end, a recent report utilized scRNA-seq to detect barcoded AAVs in mouse cortical cells^15^, but it remained challenging to detect RNA polymerase (Pol) II transcribed AAV transcripts in an efficient and cell-type unbiased way at the single-cell level.

We aimed to combine efficient RNA barcoding and STARmap PLUS detection to spatially profile cell-type-resolved tropisms of AAV in single cells (Fig. 4a). First, to enhance the accumulated copy number of RNA barcodes produced from the AAV, we designed the barcode in the form of circular RNA^19^ under a Pol III-transcribed U6 promoter (Extended Data Fig. 1c). A circular RNA lacks exposed 5’- and 3’-ends and thus exhibits extended RNA stability^45^, making it the ideal vector for RNA barcodes. Meanwhile, instead of using any Pol II promoters with potential cell-type bias^46^, we used a generic U6 promoter to optimize the RNA barcode expression across all cell types. Second, the unique *in situ* hybridization and amplification processes in STARmap PLUS allowed absolute quantification of RNA molecule copy numbers in individual cells (Extended Data Fig. 8a). Finally, measuring the cellular transcriptome and AAV RNA barcodes simultaneously in the same cell in the same tissue sample allowed cell-type-resolved quantification of AAV tropisms.

**Figure 4.**
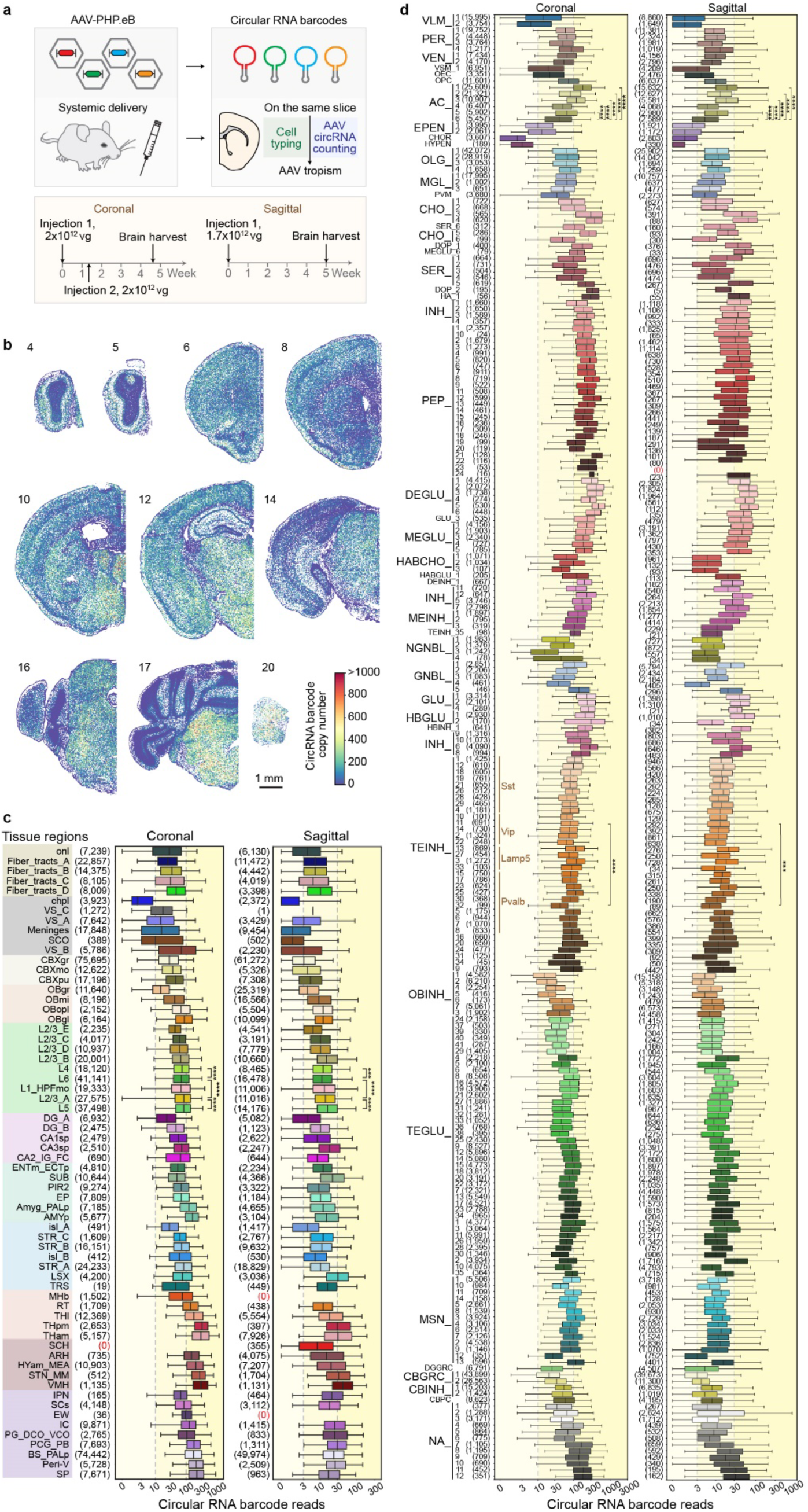
AAV-PHP.eB tropism across adult mouse CNS tissue regions and molecular cell types. **a**, Schematics of AAV-PHP.eB tropism characterization strategy across the adult mouse CNS. vg, viral genome. **b**, Circular RNA expression on representative coronal slices. Each dot represents a cell color-coded by its AAV barcode expression level. **c**,**d**, Boxplots of circular RNA expression levels across molecular tissue regions (**c**) and subcluster cell types (**d**) in coronal and sagittal slices, respectively. Molecular tissue regions were color-grouped by anatomical definitions and ranked by mean tissue circular RNA barcode expression level within the group. Boxplot elements: vertical line, median; box, first quartile to the third quartile; whiskers, 2.5-97.5%. Numbers in parentheses, number of cells in the group. Abbreviations for tissue region and cell type are the same as Figure 2 legend. *P* values were calculated by the two-sample two-tailed Kolmogorov-Smirnov test between indicated groups. \*\*\**P* < 0.001, \*\*\*\**P* < 0.0001. Data are provided in the accompanying Source Data file.

As a proof of concept, we characterized the CNS-wide, cell-type tropism of PHP.eB^5, 6^ at single- cell resolution. Coronal and sagittal sectioning samples were collected from mice injected with different doses and barcodes of PHP.eB (Fig. 4a). After *in situ* sequencing and cell segmentation, we assigned RNA barcode-derived amplicons to each cell based on the cell identities of their nearest mRNA amplicons in the 3D space (see Methods, Extended Data Fig. 8a). We observed a good correlation between the coronal and sagittal replicates (Pearson’s *r* ≥ 0.829, *P* < 0.0001, Extended Data Fig. 9a,b), supporting the potency and robustness of our experimental and computational approaches to cell-type tropism profiling.

We first assessed AAV-PHP.eB tropism across molecularly defined tissue regions. Among all brain regions, we observed generally higher RNA barcode expression in the brain stem than cerebrum (Fig. 4b, Extended Data Fig. 8b). Notably, this result is different from the previously reported protein-based readout^5^, where GFP fluorescence signals controlled by the CAG promoter were higher in the cerebrum than the brain stem. Such differences suggest that U6- based Pol III and CAG-based Pol II transcription activities may vary differently across cell types and tissue regions. Further examining the tropism across molecular tissue regions (Fig. 4c) revealed that circular RNA expression is in general lower in glia-rich regions (e.g., fiber tracts, ventricles, meninges, choroid plexus, and subcommissural organ) than in neuron-rich regions. Among those neuron-rich regions, we further noticed that molecular regions for granular cells, including CBXgr (cerebellar cortex granular layer), OBgr (olfactory bulb granular layer), DG_A (dentate gyrus granular layer), and isl_A (islands of Calleja), tend to have lower RNA barcode levels than agranular areas (e.g., cerebral cortex). Next, the cerebral regions showed modest AAV transduction, and the expression level of RNA barcodes did not vary dramatically across cortical layers despite a slight enrichment at L5 followed by L2/3_A (Fig. 4c). Lastly, tissue regions in the thalamus, hypothalamus, midbrain, and hindbrain exhibit the highest barcode levels (Fig. 4c).

Next, we asked how AAV-PHP.eB tropism varies across molecular cell types. In addition to recapitulating the known tropism of PHP.eB towards neurons and astrocytes^5, 15^ (Extended Data Fig. 9c), we uncovered new viral tropism features across fine molecular cell types at unprecedented resolution. Among astrocytes, we confirmed the preference of PHP.eB for *Myoc*^-^ astrocytes (AC_1∼5) over *Myoc*^+^ ones (AC_6)^15^ (*P* < 0.0001, two-sided Kolmogorov-Smirnov test; Fig. 4d, Extended Data Fig. 3a). In other glial cells, oligodendrocytes and oligodendrocyte precursor cells show modest PHP.eB transduction, followed by immune, vascular, and olfactory ensheathing cells. Epithelial cells are the lowest among all cell types in RNA barcode expression, including EPEN, CHOR, and SCO hypendymal cells (HYPEN) (Extended Data Fig. 9c). Within neurons, TEGLU and TEINH neurons in the cerebral cortex show similar PHP.eB transduction levels (Extended Data Fig. 9c). In contrast, excitatory neurons have higher RNA barcode expression than inhibitory ones in the di- and mesencephalon (*P* < 0.0001, two-sided Kolmogorov-Smirnov test; Extended Data Fig. 9c). We further characterized PHP.eB tropism profiles among subclusters (Fig. 4d). To highlight a few, we found that cholinergic and monoaminergic neurons, di- and mesencephalon excitatory neurons, and di- and mesencephalon inhibitory neurons showed large variation in PHP.eB transduction among their respective subtypes (Extended Data Fig. 9d, Fig. 4d). As another example, the PHP.eB tropisms among different categories of TEINH marked by *Sst*, *Pvalb*, *Vip*, and *Lamp5* in general matched with the previously reported trend^15^: *Lamp5^+^*> *Pvalb^+^* ∼ *Sst^+^* > *Vip^+^* (Extended Data Fig. 9e). However, our data further illustrated that this trend was not necessarily true at the subcluster level (e.g., *Vip^+^* TEINH_11 > *Pvalb^+^* TEINH_32, *P* < 0.001, two-sided Kolmogorov-Smirnov test; Fig. 4d). In total, deep cell typing at the subtype level is critical for achieving accurate profiling of AAV tropisms.

## Discussion

This work presents a comprehensive spatial atlas across the whole mouse CNS at 200 nm resolution, encompassing over one million cells with 1,022 genes measured by STARmap PLUS. We clustered and annotated 26 main cell types, 231 subtypes, and 64 molecular tissue regions in 3D space with ClusterMap (Fig. 1-2), providing a road map for investigating CNS-wide gene-expression patterns and cell-type diagrams in the context of brain anatomy. Leveraging existing mouse CNS scRNA-seq data, we predicted transcriptome profiles of individual cells by data integration and gene imputation (Fig. 3). In addition to the spatial cell typing, we demonstrated, for the first time, the quantitative tissue-region- and cell-type-resolved tropisms of a whole-brain gene delivery tool, AAV-PHP.eB (Fig. 4). We complemented our atlas with an online database, mCNS_atlas, with exploratory interfaces (http://brain.spatial-atlas.net), serving as an open resource for neurobiological studies across molecular, cellular, and tissue levels.

Our strategy and the resulting datasets have the following advantages. First, measuring RNA molecules *in situ* minimized the disturbance from sample preparation on single-cell expression profiles, exemplified by the percentage of activated microglia in the whole microglia population (*Ccl3*^+^ or *Ccl4*^+^, 8.8% in the current atlas versus 24.6% in the scRNA-seq atlas^1^). Second, spatial profiling of molecular cell types further enabled molecular tissue segmentation and molecule-based data integration among different samples and different technology platforms, generating a unified molecular definition of tissue regions that may be more accurate and reproducible than human-annotated tissue anatomy. Finally, multiplexing measurements in the same sample allowed experimental integration of endogenous cellular features with exogenously introduced genetic labeling or perturbation, as illustrated here by the tropism characterization of PHP.eB in the mouse CNS (Fig. 4a). Importantly, this systematic strategy can be readily adapted to simultaneously profile tropisms of multiple AAV capsid variants or screen various cell-type-specific promoter and enhancer sequences within the same sample by barcoding each variant, enabling cell-type resolved, tissue-level characterization of therapeutics distribution and responses^47^. Such characterization can serve as a screening platform and selection guide for AAV variants to maximize transgene expression in targeted cell types in research and therapeutic applications.

In conclusion, we provided large-scale, organ-wide, single-cell, and spatially resolved transcriptome profiles of the mouse CNS at molecular resolution. We envision the opportunities to integrate our datasets with additional modalities including chromatin measurements, cell morphology, and cell-cell communications^48, 49^. The scalable experimental and computational frameworks demonstrated here can be readily applied to map whole-organ and whole-animal cell atlases across species and disease models to study development, evolution, and disorders.

## Data Availability statement

The STARmap PLUS sequencing data of this study are available on the Single Cell Portal (https://singlecell.broadinstitute.org/single_cell/study/SCP1830). We also introduced an interactive online database (http://brain.spatial-atlas.net) for exploratory analysis and hypothesis generation.

## Code Availability statement

The following packages^50–63^ and software were used in the data analysis: ClusterMap is implemented based on MATLAB R2019b and Python 3.6. The following packages and software were used in data analysis: UCSF ChimeraX 1.0, ImageJ 1.51, MATLAB R2019b, R 4.0.4, Rstudio 1.4.1106, Jupyter Notebook 6.0.3, Anaconda 2-2-.02, h5py 3.1.0, hdbscan 0.8.36, hdf5 1.10.4, matplotlib 3.1.3, seaborn 0.11.0, scanpy 1.6.0, numpy 1.19.4, scipy 1.6.3, pandas 1.2.3, scikit-learn 0.22, umap-learn0.4.3, pip 21.0.1, numba 0.51.2, tifffile 2020.10.1, scikit-image 0.18.1, itertools 8.0.0. The code that supports the analyses in this study is available at https://github.com/wanglab-broad/mCNS-atlas.

## Acknowledgments

We thank Dr. Ken Y. Chan from Ben Deverman’s lab in the Stanley Center for Psychiatric Research at the Broad Institute of MIT and Harvard for guidance on AAV packaging and delivery, Dr. Ming Pan from Nicol’s lab at the Broad Institute of MIT and Harvard for technical assistance, Kamal Maher from Wang lab for valuable input in tissue region identification, and Jane Salant from Liu lab for manuscript editing. X.W. acknowledges the support from the Searle Scholars Program, Thomas D. and Virginia W. Cabot Professorship, Edward Scolnick Professorship, Ono Pharma Breakthrough Science Initiative Award, and NIH DP2 New Innovator Award. J.L. acknowledges the support from the Aramont Fund. H.S. is supported by Helen Hay Whitney Foundation Postdoctoral Fellowship. Y.H. is supported by the James Mills Peirce Fellowship from the Graduate School of Arts and Sciences of Harvard University.

## Author contributions

X.W. and H.S. designed the project. Y.Z. packaged the rAAV. Y.Z. and P.T. performed animal experiments. H.S. and Y.Z. performed the STARmap PLUS data acquisition. Y.H, J.H, B.W, H.S., and Y.Z. analyzed the data. Z.T. and M.W. implemented the online data portal. Z.L., A.L., J.R., and Y.T. assisted with experiments. X.T. helped in computation pipeline optimization. H.S., Y.H., and B.W. prepared figures. H.S., Y.H., Y.Z., J.H., B.W., J.L., and X.W. wrote the manuscript with inputs from all authors. X.W. supervised the study.

## Additional information

Correspondence and requests for the material should be addressed to Xiao Wang.

## Supplementary information

**Supplementary Table 1**: 1,022 gene list of STARmap PLUS and the 5-nt code for each gene.

**Supplementary Table 2**: Sequences for SNAIL probes and SEDAL seq probes.

**Supplementary Table 3**: Detailed information for mice and tissue slices in the atlas.

**Supplementary Table 4**: Subcluster cell-type annotation, including cell type symbols, cell numbers, color codes, cell type description, and spatial distribution.

**Supplementary Table 5**: Tissue region annotation, including tissue region symbol, description, featured cell types, and marker genes.

**Supplementary Table 6**: Immediate mapping performance evaluation, tissue-region mean expression of imputed genes on the sample slice 12, and putative marker genes of medial habenula on the sample slice 12.

---

**Extended Data Figure 1.**
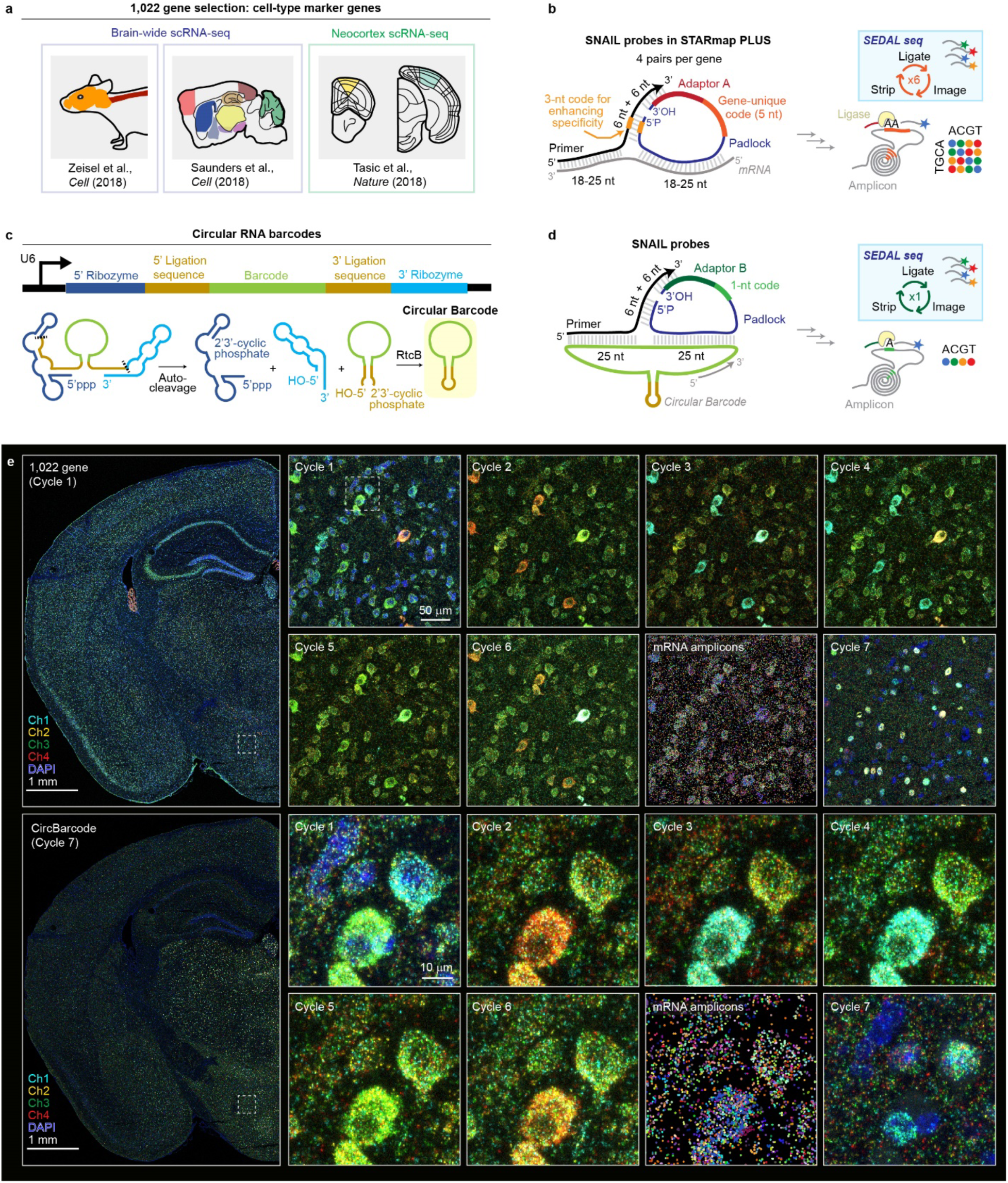
Probe designs and raw fluorescent images of adult mouse CNS STARmap PLUS. **a**, Mouse brain single-cell RNA-seq (scRNA-seq) sources for STARmap PLUS 1,022 gene-list selection. **b**, SNAIL probes (primer and padlock probes) for 1,022 endogenous genes. The padlock probe contains a 5-nt gene-unique identifier, which is amplified during rolling- cycle amplification and read out by six cycles of sequential SEDAL seq with adaptor sequence *A*. **c**, Schematics showing the construct design and biogenesis of circular RNA barcodes. **d**, SNAIL probes for circular RNA barcodes. Each barcode is converted to a 1-nt identifier and read out by one additional cycle of SEDAL seq with adaptor sequence *B*. **e**, Raw fluorescent images of SEDAL seq of brain slice 12. Left panels show the image stack maximum projection of SEDAL seq cycles 1 and 7, merged into an entire half slice. The top-right panels show zoomed-in views of SEDAL seq cycles 1 to 7 and amplicons colored by gene identity from the square highlighted in the left panels. The bottom-right panels show zoomed-in views of the square highlighted in the top right panels.

**Extended Data Figure 2.**
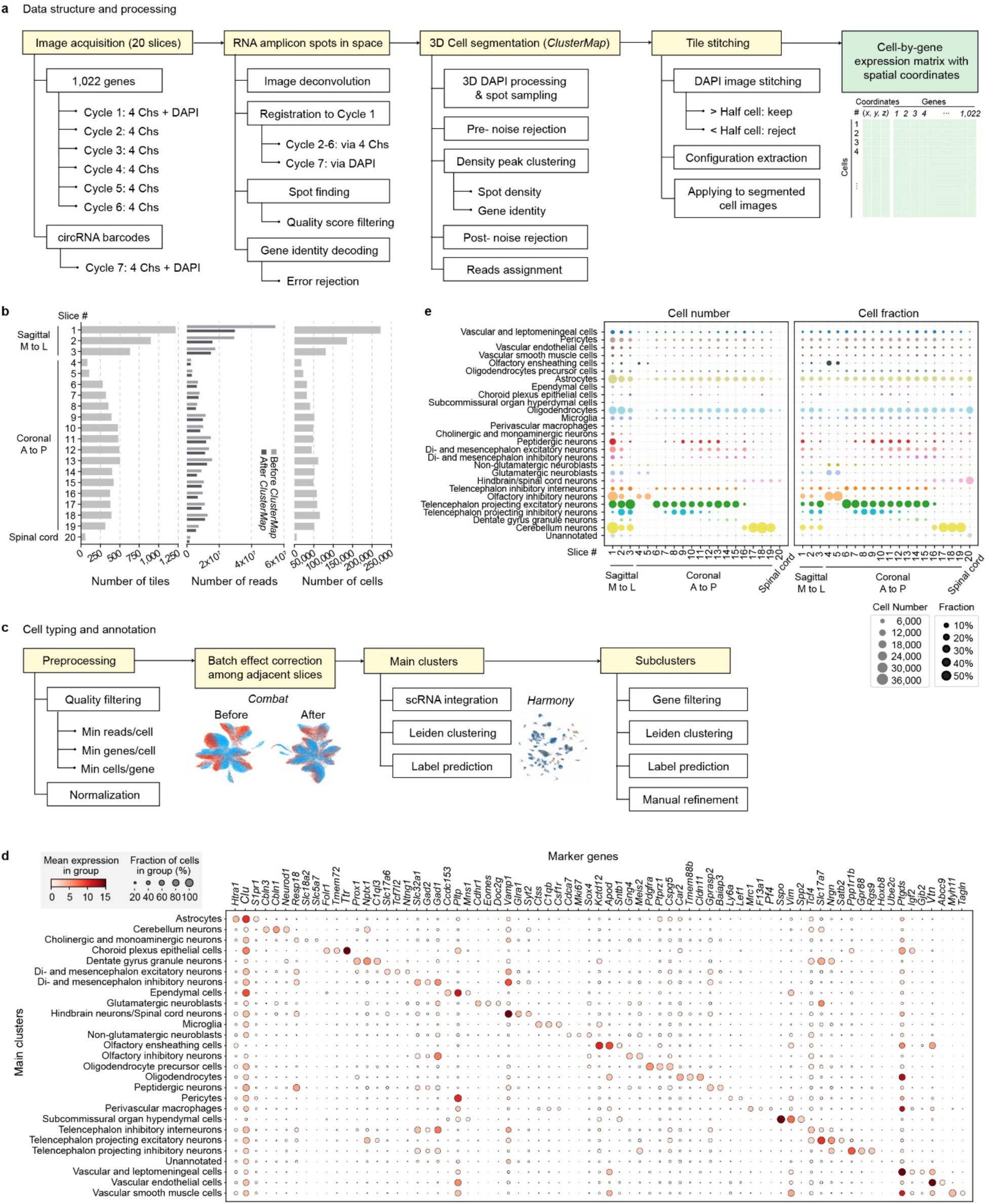
Spatial cell typing workflow and data quality. **a**, Data structure of the study and the workflow from raw images to a cell-by-gene matrix with cell spatial coordinates. Chs, channels. **b**, Summary of the number of tiles (i.e., imaging area), reads, and cells in each tissue sample slice. M, medial; L, lateral; A, anterior; P, posterior. **c**, Workflow of cell typing and annotation. **d**, Dotplots of the top three marker genes for each main cluster. **e**, Main-cluster cell- type composition of each tissue sample slice. Left, absolute cell number; right, cell fraction normalized within each tissue slice. Data are provided in the accompanying Source Data file.

**Extended Data Figure 3.**
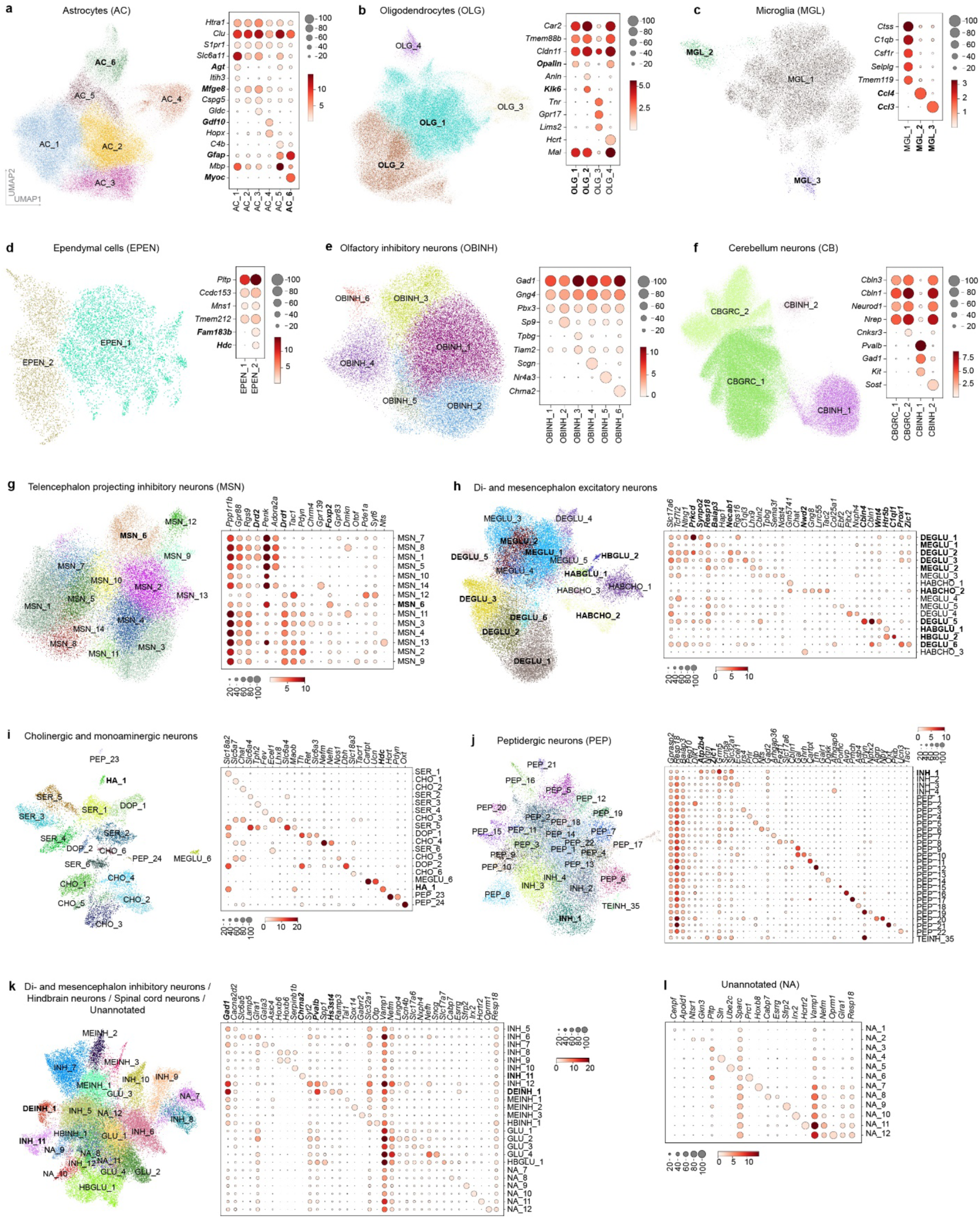
Subclustering of main cell types. **a**-**k**, Uniform Manifold Approximation and Projection (UMAP) maps (left) and marker gene dotplots (right) of main clusters colored by cell subcluster identities, for astrocytes (AC, **a**), oligodendrocytes (OLG, **b**), microglia (MGL, **c**), ependymal cells (EPEN, **d**), olfactory inhibitory neurons (OBINH, **e**), cerebellum neurons (CB, **f**), telencephalon projecting inhibitory neurons (MSN, **g**), di- and mesencephalon excitatory neurons (**h**), cholinergic and monoaminergic neurons (**i**), peptidergic neurons (PEP or INH, **j**), and di- and mesencephalon inhibitory neurons / hindbrain neurons/ spinal neurons / unannotated (**k**). **l**, Marker gene dotplot for unannotated (NA) clusters. Dot sizes, fraction of cells in the group; color bars, mean expression level in the group. Cell types and genes mentioned in the main text are bolded. Also see Methods and Supplementary Table 4.

**Extended Data Figure 4.**
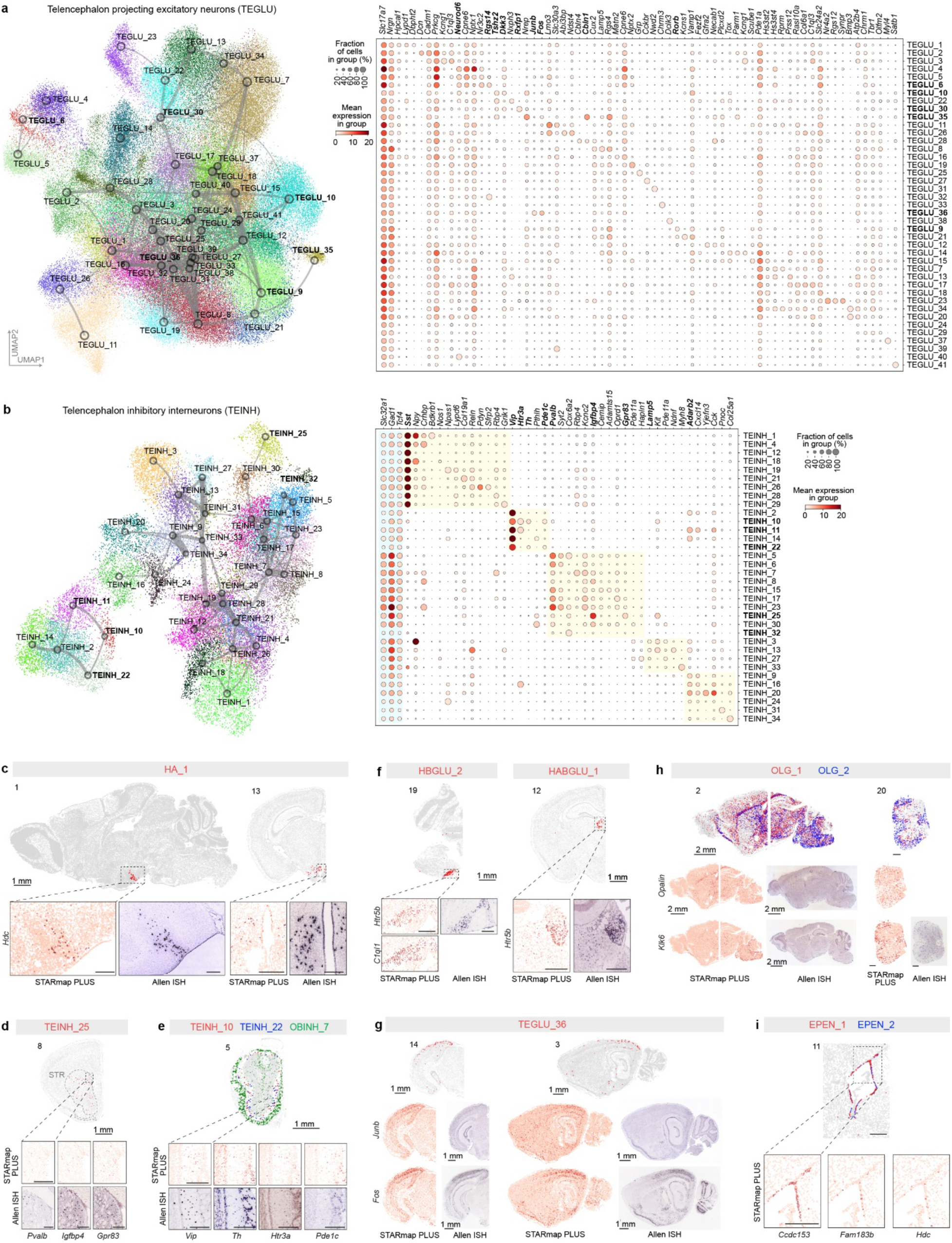
Subclustering of telencephalon neurons and spatial maps of representative subcluster cell types. **a**,**b**, Overlapped UMAP map and constellation plot^31^ of main clusters colored by cell subcluster identities (left) and marker gene dotplots (right), for telencephalon projecting excitatory neurons (TEGLU, **a**) and telencephalon inhibitory interneurons (TEINH, **b**). Cell types and marker genes mentioned in the main text are bolded. Also see Methods and Supplementary Table 4. **c**-**i**, Cell-type spatial maps, (zoomed-in) spatial expression heatmap of cell-type marker genes measured by STARmap PLUS, and corresponding *In Situ* Hybridization images of the marker genes from the Allen Mouse Brain ISH database^27^, for subcluster cell types HA_1 (**c**), TEINH_25 (**d**), TEINH_10, TEINH_22, and OBINH_7 (**e**), HBGLU_2 and HABGLU_1 (**f**), TEGLU_36 (**g**), OLG_1 and OLG_2 (**h**), and EPEN_1 and EPEN_2 (**i**). Each dot represents a cell color-coded by its cell-type symbol. Scale bars, 250 μm if not indicated. HA, histaminergic neurons; OBINH, olfactory inhibitory neurons; HBGLU, hindbrain excitatory neurons; HABGLU, habenular excitatory neurons; OLG, oligodendrocytes; EPEN, ependymal cells; STR, striatum.

**Extended Data Figure 5.**
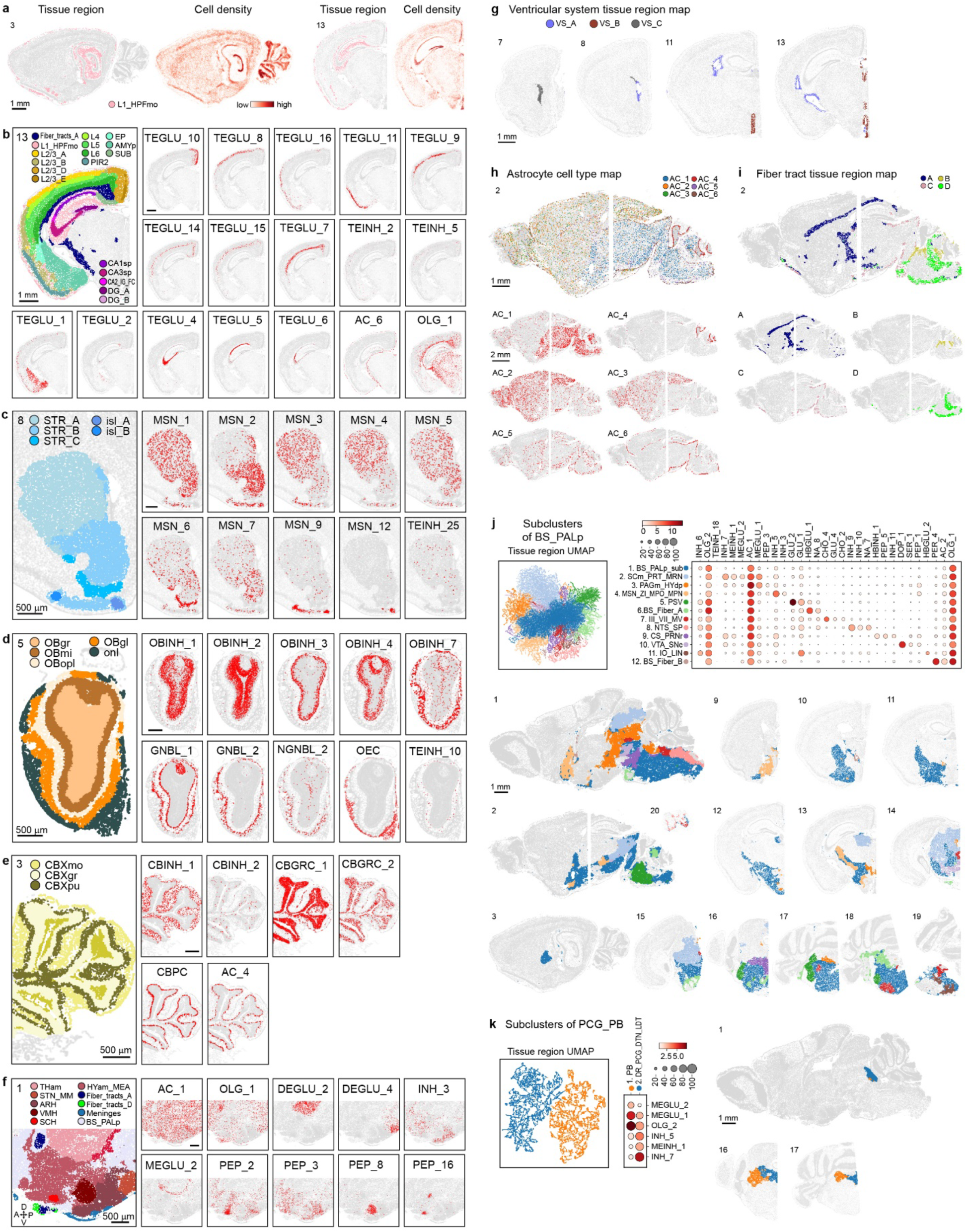
Cell density, marker cell types, and finer clustering of molecular tissue regions. **a**, Representative maps of the molecular tissue region L1_HPFmo and the corresponding spatial heatmaps colored by relative cell densities. Each dot represents a cell. **b**- **f**, Representative tissue region organizations and spatial maps of relevant marker cell types in the cerebral cortex (**b**), striatum (**c**), olfactory bulb (**d**), cerebellum (**e**), and medial hypothalamus (**f**). Each dot represents a cell colored by its molecular tissue region identity. A, anterior; P, posterior; D, dorsal; V, ventral. **g**, Molecular tissue region maps for ventricular systems. **h**, Overlaid and individual spatial maps of astrocyte subtypes in sample slice 2. **i**, Overlaid and individual molecular tissue region maps for fiber tracts in sample slice 2. **j**,**k**, Neighborhood cell- type composition (NCC) UMAPs, marker cell-type dotplots, and spatial molecular tissue region maps for subclusters in BS_PALp (**j**) and PCG_PB (**k**). Abbreviations for tissue region and cell type: SCm, superior colliculus, motor-related; PRT, pretectal region; MRN, midbrain reticular nucleus; PAGm, medial periaqueductal gray; HYdp, dorsal-posterior hypothalamus; MSC, medial septal complex; ZI, zona incerta; MPO, medial preoptic area; MPN, medial preoptic nucleus; PSV, principal sensory nucleus of the trigeminal; III, oculomotor nucleus; VII, facial motor nucleus; MV, medial vestibular nucleus; NTS, nucleus of the solitary tract; SP, spinal cord related; CS, superior central nucleus raphe; PRNr, pontine reticular nucleus; VTA, ventral tegmental area; SNc, substantia nigra, compact part; IO, inferior olivary complex; LIN, linear nucleus of the medulla. Also see Figure 2 legend.

**Extended Data Figure 6.**
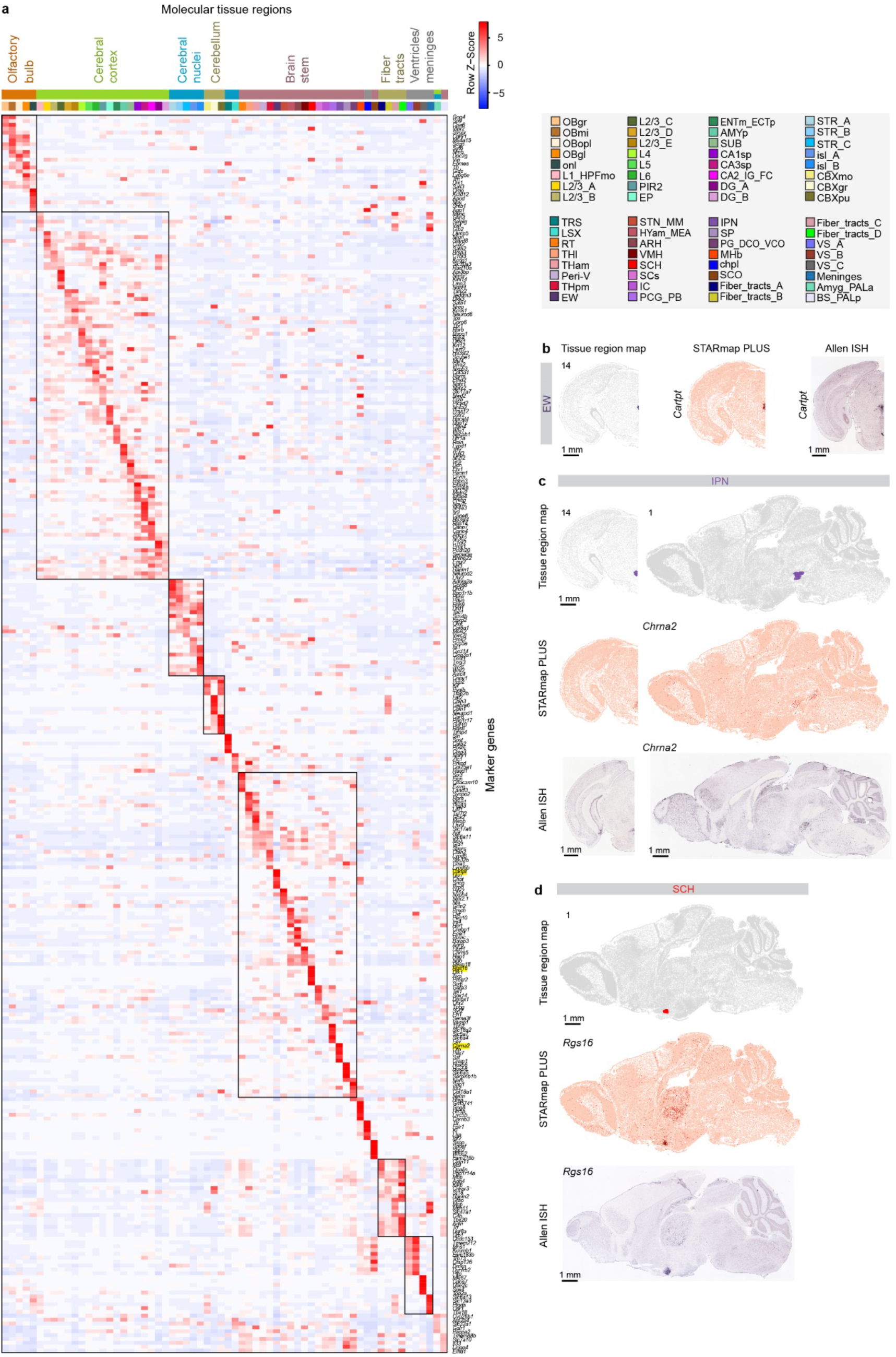
Marker genes of molecular tissue regions. **a**, Expression heatmap of the top five marker genes for each tissue region. The mean expression level of a gene across all cells in a tissue region was calculated and then scaled across tissue regions in the heatmap. Marker genes visualized in **b**-**d** were highlighted in yellow rectangles. **b**-**d**, Spatial tissue region maps, spatial expression heatmaps of the tissue region marker gene measured by STARmap PLUS, and *In Situ* Hybridization images of the marker gene from the Allen Mouse Brain ISH database^27^, for molecular tissue regions Edinger-Westphal nucleus (EW, **b**), interpeduncular nucleus (IPN, **c**), and hypothalamic suprachiasmatic nucleus (SCH, **d**). Abbreviations of tissue regions were based on Allen Mouse Brain Reference Atlas^24^. Also see Figure 2 legend. Data are provided in the accompanying Source Data file.

**Extended Data Figure 7.**
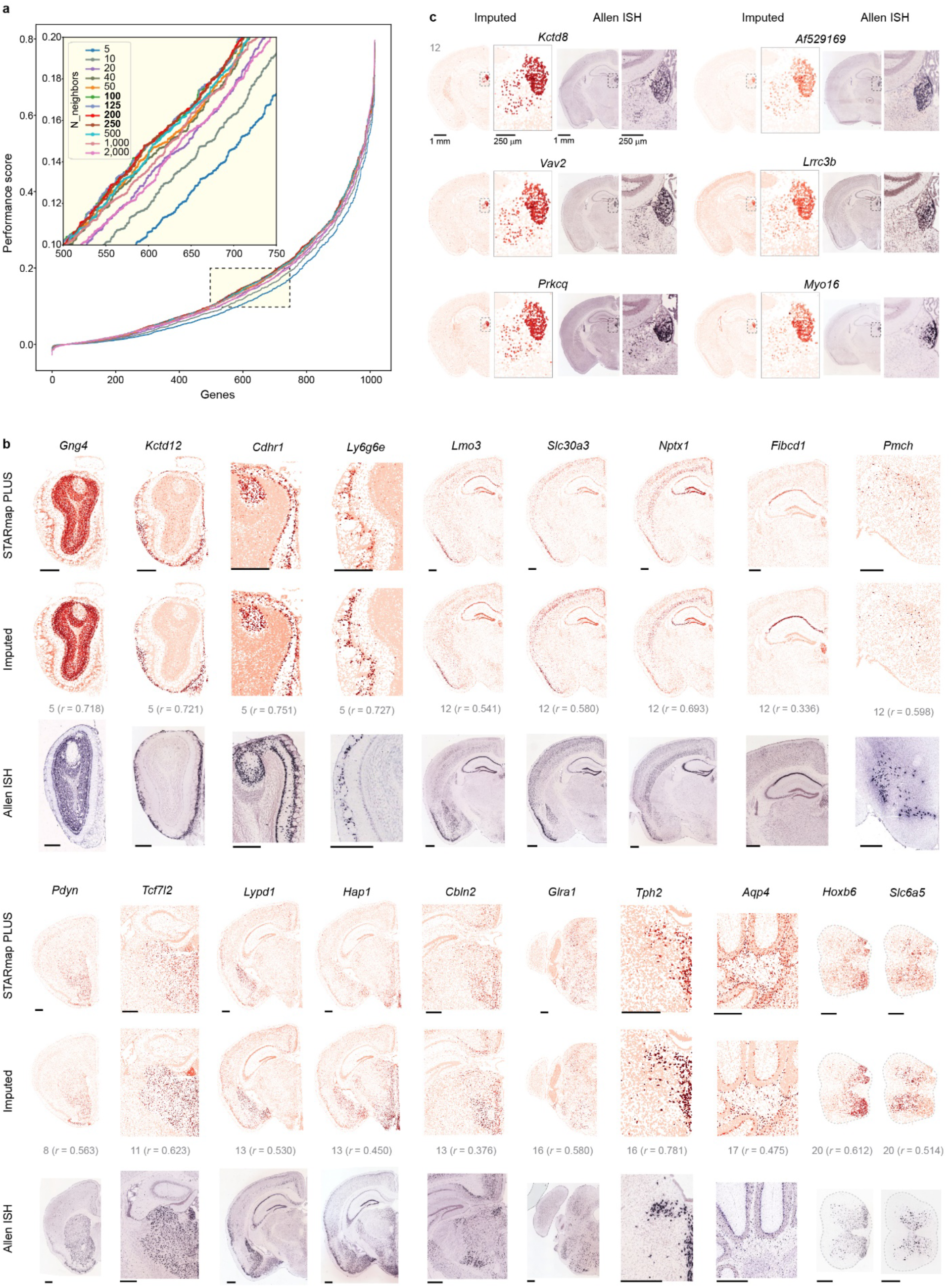
Imputation parameter optimization and performance evaluation. **a**, Cumulative curves of the performance scores across 1,022 genes in the immediate mapping using different numbers of scRNA-seq atlas cell nearest neighbors. The upper-left inset shows a zoomed-in view of the rectangular region highlighted in the bottom right. **b**, More examples of the comparison of imputed spatial gene expression with measured expression from STARmap PLUS and Allen Mouse Brain *In Situ* Hybridization (ISH) database^27^. Each dot represents a cell colored by the expression level of a specified mRNA. Scale bar, 0.5 mm. The sample slice number and Pearson correlation coefficient *r* between the STARmap PLUS measurements and the imputed results were labeled in gray. **c**, Imputed spatial expression heatmap of putative medial habenula marker genes and paired ISH images from the Allen Mouse Brain ISH database^27^. Data are provided in the accompanying Source Data file.

**Extended Data Figure 8.**
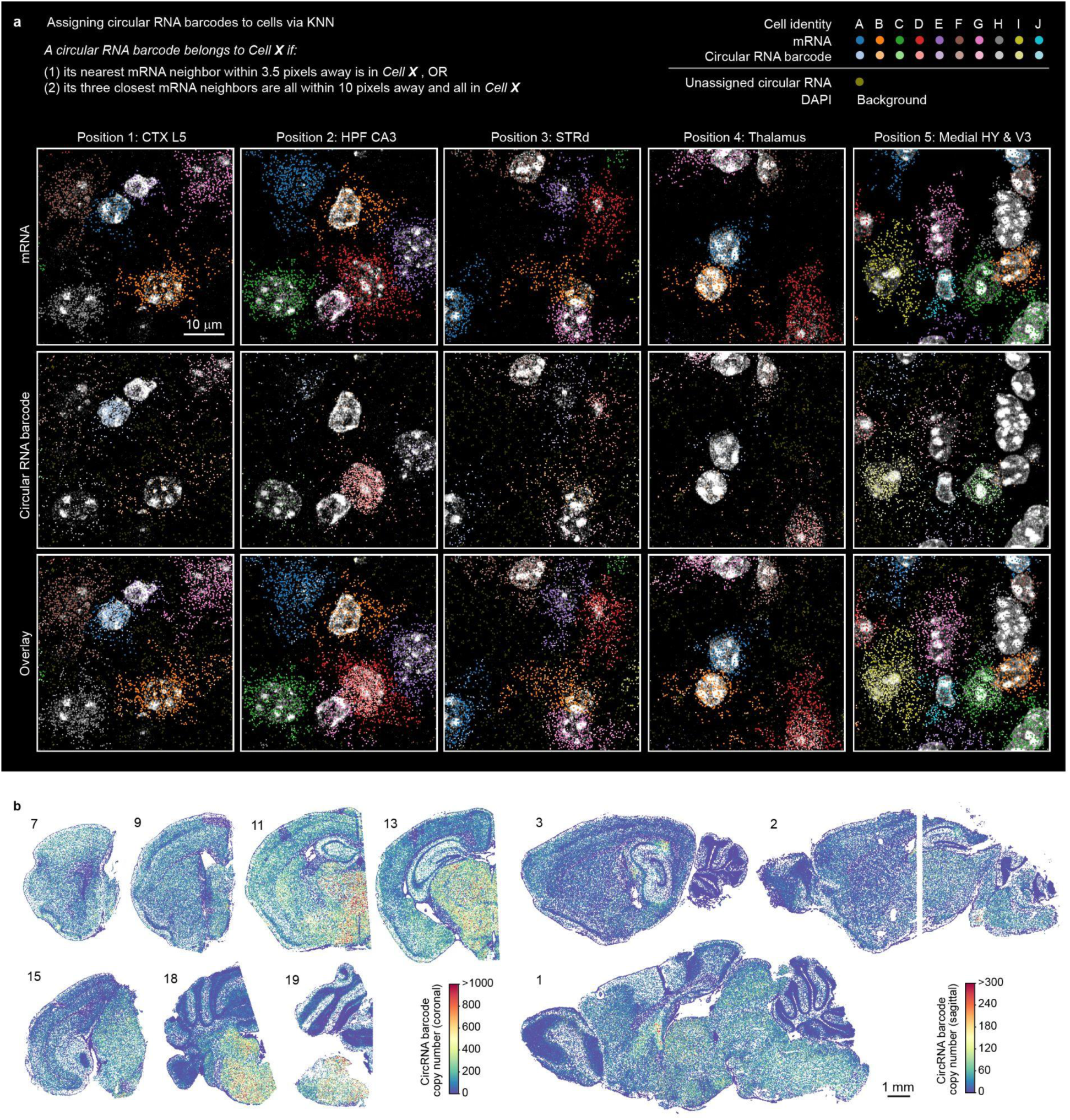
Circular RNA barcode assignment and spatial distribution. **a**, Examples illustrating how circular RNA barcodes are assigned into individual ClusterMap- identified cells. 1 pixel = 194 nm. CTX, cerebral cortex; HPF, hippocampal formation; STRd, dorsal striatum; HY, hypothalamus; V3, ventricle 3. **b**, Circular RNA expression on the rest of the coronal slices and sagittal slices. Each dot represents a cell color-coded by its barcode expression level.

**Extended Data Figure 9.**
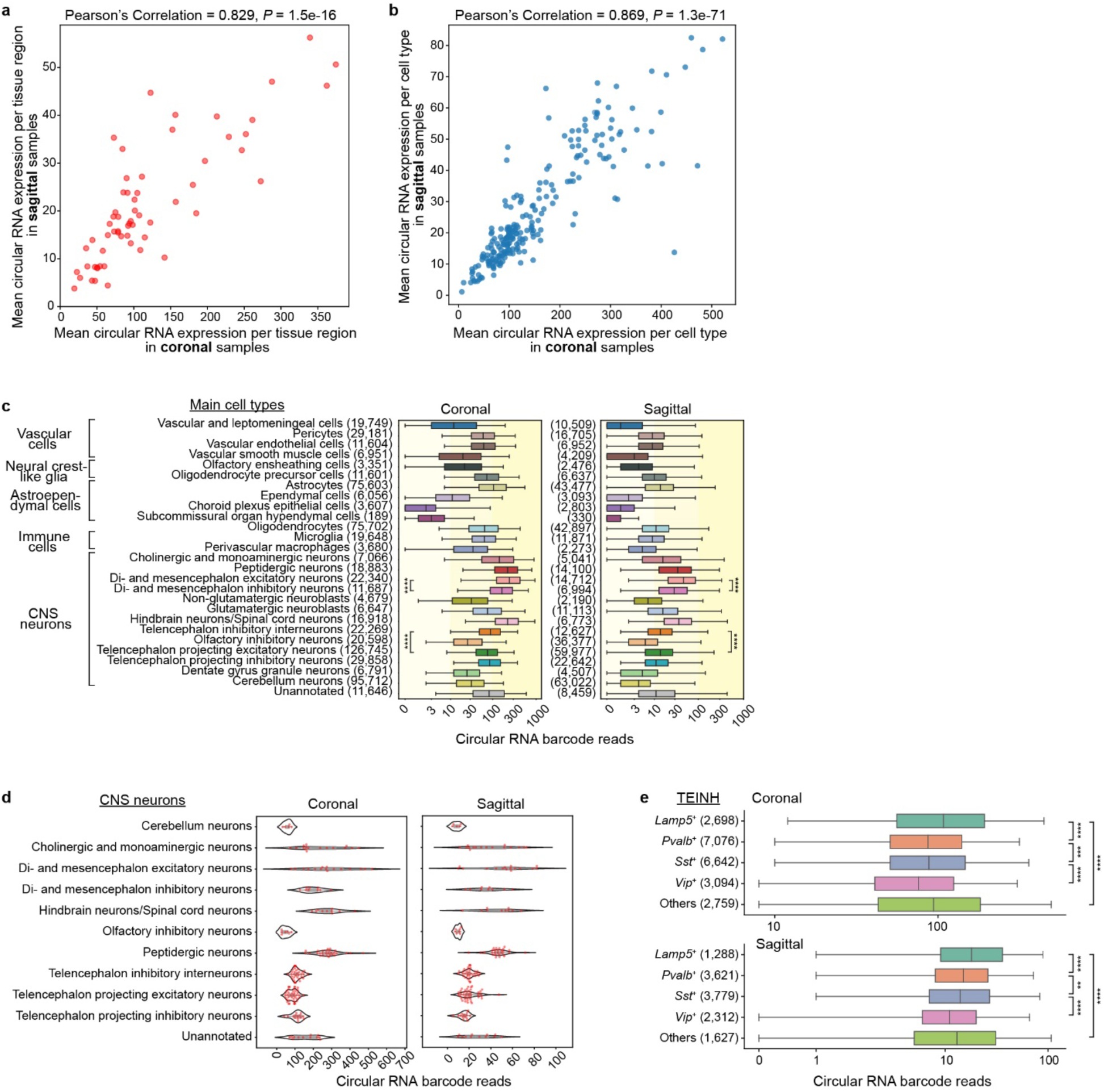
Circular RNA barcode expression in molecular tissue regions and molecular cell types. **a**,**b**, Correlation of mean circular barcode expression between coronal and sagittal replicates across molecular tissue regions (**a**) and subcluster cell types (**b**). Each dot represents a molecular tissue region (**a**) or a subcluster cell type (**b**). **c**, Circular RNA expression level across main cluster cell types. Boxplot elements: the vertical line, median; the box, first to third quartiles; whiskers, 2.5-97.5%. Numbers in parentheses, number of cells in the group. **d**, Violin plots of the mean circular RNA barcode expression level of subcluster cell types in each main neuronal cluster. Each dot represents a subcluster cell type. Boxplot elements inside the violin: the gray dot, median; the box, interquartile range; whiskers,1.5x interquartile range. **e**, Circular RNA expression level across telencephalon inhibitory interneurons (TEINH) expressing different neuronal peptides. Boxplot elements: the vertical line, median; the box, first to third quartiles; whiskers, 2.5-97.5%. Numbers in parentheses, number of cells in the group. *P* values were calculated by two-sample two-tailed Kolmogorov-Smirnov test between indicated groups (**c**,**e**). \*\**P* < 0.01, \*\*\**P* < 0.001, \*\*\*\**P* < 0.0001. Data are provided in the accompanying Source Data file.

## References

1. Zeisel, A. et al. Molecular Architecture of the Mouse Nervous System. Cell 174, 999–1014.e22 (2018).

2. Saunders, A. et al. Molecular Diversity and Specializations among the Cells of the Adult Mouse Brain. Cell 174, 1015–1030.e16 (2018).

3. Wang, X. et al. Three-dimensional intact-tissue sequencing of single-cell transcriptional states. Science vol. 361 (2018).

4. Hu Zeng, Jiahao Huang, Haowen Zhou, William J. Meilandt, Borislav Dejanovic, Yiming Zhou, Christopher J. Bohlen, Seung-Hye Lee, Jingyi Ren, Albert Liu, Hao Sheng, Jia Liu, Morgan Sheng, Xiao Wang. Integrative in situ mapping of single-cell transcriptional states and tissue histopathology in an Alzheimer’s disease model. bioRxiv (2022) doi:10.1101/2022.01.14.476072.

5. Chan, K. Y. et al. Engineered AAVs for efficient noninvasive gene delivery to the central and peripheral nervous systems. Nature Neuroscience vol. 20 1172–1179 (2017).

6. Goertsen, D. et al. AAV capsid variants with brain-wide transgene expression and decreased liver targeting after intravenous delivery in mouse and marmoset. Nat. Neurosci. 25, 106–115 (2022).

7. Ortiz, C., Carlén, M. & Meletis, K. Spatial Transcriptomics: Molecular Maps of the Mammalian Brain. Annu. Rev. Neurosci. 44, 547–562 (2021).

8. Rao, A., Barkley, D., França, G. S. & Yanai, I. Exploring tissue architecture using spatial transcriptomics. Nature 596, 211–220 (2021).

9. Liao, J., Lu, X., Shao, X., Zhu, L. & Fan, X. Uncovering an Organ’s Molecular Architecture at Single-Cell Resolution by Spatially Resolved Transcriptomics. Trends Biotechnol. 39, 43–58 (2021).

10. Ortiz, C. et al. Molecular atlas of the adult mouse brain. Sci Adv 6, eabb3446 (2020).

11. Moffitt, J. R. et al. Molecular, spatial, and functional single-cell profiling of the hypothalamic preoptic region. Science 362, (2018).

12. Chen, R. et al. Decoding molecular and cellular heterogeneity of mouse nucleus accumbens. Nat. Neurosci. 24, 1757–1771 (2021).

13. Zhang, M. et al. Spatially resolved cell atlas of the mouse primary motor cortex by MERFISH. Nature 598, 137–143 (2021).

14. Palfi, A., et al. AAV-PHP.eB transduces both the inner and outer retina with high efficacy in mice. Mol Ther Methods Clin Dev 25, 236–249 (2022).

15. Brown, D. et al. Deep Parallel Characterization of AAV Tropism and AAV-Mediated Transcriptional Changes Single-Cell RNA Sequencing. Front. Immunol. 12, 730825 (2021).

16. He, Y. et al. ClusterMap for multi-scale clustering analysis of spatial gene expression. Nat. Commun. 12, 5909 (2021).

17. Tasic, B. et al. Shared and distinct transcriptomic cell types across neocortical areas. Nature 563, 72–78 (2018).

18. Xu, Q., Schlabach, M. R., Hannon, G. J. & Elledge, S. J. Design of 240,000 orthogonal 25mer DNA barcode probes. Proc. Natl. Acad. Sci. U. S. A. 106, 2289–2294 (2009).

19. Litke, J. L. & Jaffrey, S. R. Highly efficient expression of circular RNA aptamers in cells using autocatalytic transcripts. Nat. Biotechnol. 37, 667–675 (2019).

20. Johnson, W. E., Li, C. & Rabinovic, A. Adjusting batch effects in microarray expression data using empirical Bayes methods. Biostatistics 8, 118–127 (2007).

21. Korsunsky, I. et al. Fast, sensitive and accurate integration of single-cell data with Harmony. Nat. Methods 16, 1289–1296 (2019).

22. Traag, V. A., Waltman, L. & van Eck, N. J. From Louvain to Leiden: guaranteeing well-connected communities. Sci. Rep. 9, 5233 (2019).

23. Fujita, A. et al. Hypothalamic Tuberomammillary Nucleus Neurons: Electrophysiological Diversity and Essential Role in Arousal Stability. J. Neurosci. 37, 9574–9592 (2017).

24. Allen reference atlases :: Atlas viewer. http://atlas.brain-map.org.

25. Bao, F. et al. Integrative spatial analysis of cell morphologies and transcriptional states with MUSE. Nat. Biotechnol. 1–10 (2022).

26. Tepe, B. et al. Single-Cell RNA-Seq of Mouse Olfactory Bulb Reveals Cellular Heterogeneity and Activity-Dependent Molecular Census of Adult-Born Neurons. Cell Rep. 25, 2689 (2018).

27. Lein, E. S. et al. Genome-wide atlas of gene expression in the adult mouse brain. Nature 445, 168–176 (2007).

28. Wang, Q. et al. The Allen Mouse Brain Common Coordinate Framework: A 3D Reference Atlas. Cell 181, 936–953.e20 (2020).

29. Peters, A. AP_histology. https://github.com/petersaj/AP_histology (2019).

30. Philip Shamash, Matteo Carandini, Kenneth Harris, Nick Steinmetz. A tool for analyzing electrode tracks from slice histology. bioRxiv (2018) doi:10.1101/447995.

31. Yao, Z. et al. A taxonomy of transcriptomic cell types across the isocortex and hippocampal formation. Cell 184, 3222–3241.e26 (2021).

32. Yamawaki, N., Borges, K., Suter, B. A., Harris, K. D. & Shepherd, G. M. G. A genuine layer 4 in motor cortex with prototypical synaptic circuit connectivity. Elife 3, e05422 (2014).

33. Urbán, N. & Guillemot, F. Neurogenesis in the embryonic and adult brain: same regulators, different roles. Front. Cell. Neurosci. 8, 396 (2014).

34. Sanders, M., Petrasch-Parwez, E., Habbes, H.-W., Düring, M. V. & Förster, E. Postnatal Developmental Expression Profile Classifies the Indusium Griseum as a Distinct Subfield of the Hippocampal Formation. Front Cell Dev Biol 8, 615571 (2020).

35. Park, S.-B. et al. The fasciola cinereum subregion of the hippocampus is important for the acquisition of visual contextual memory. Prog. Neurobiol. 210, 102217 (2022).

36. Lim, D. A. & Alvarez-Buylla, A. The Adult Ventricular-Subventricular Zone (V-SVZ) and Olfactory Bulb (OB) Neurogenesis. Cold Spring Harb. Perspect. Biol. 8, (2016).

37. Lars E. Borm, Alejandro Mossi Albiach, Camiel C.A. Mannens, Jokubas Janusauskas, Ceren Özgün, David Fernández-García, Rebecca Hodge, Ed S. Lein, Simone Codeluppi, Sten Linnarsson. Scalable in situ single-cell profiling by electrophoretic capture of mRNA. bioRxiv (2022) doi:10.1101/2022.01.12.476082.

38. Topilko, T. et al. Edinger-Westphal peptidergic neurons enable maternal preparatory nesting. Neuron (2022) doi:10.1016/j.neuron.2022.01.012.

39. Lotfipour, S. et al. Targeted deletion of the mouse α2 nicotinic acetylcholine receptor subunit gene (Chrna2) potentiates nicotine-modulated behaviors. J. Neurosci. 33, 7728– 7741 (2013).

40. Doi, M. et al. Circadian regulation of intracellular G-protein signalling mediates intercellular synchrony and rhythmicity in the suprachiasmatic nucleus. Nat. Commun. 2, 327 (2011).

41. Lohoff, T. et al. Integration of spatial and single-cell transcriptomic data elucidates mouse organogenesis. Nat. Biotechnol. 40, 74–85 (2022).

42. Wallace, M. L. et al. Anatomical and single-cell transcriptional profiling of the murine habenular complex. Elife 9, (2020).

43. Wang, D., Tai, P. W. L. & Gao, G. Adeno-associated virus vector as a platform for gene therapy delivery. Nat. Rev. Drug Discov. 18, 358–378 (2019).

44. Van Vliet, K. M., Blouin, V., Brument, N., Agbandje-McKenna, M. & Snyder, R. O. The role of the adeno-associated virus capsid in gene transfer. Methods Mol. Biol. 437, 51–91 (2008).

45. Holdt, L. M., Kohlmaier, A. & Teupser, D. Circular RNAs as Therapeutic Agents and Targets. Front. Physiol. 9, 1262 (2018).

46. Qin, J. Y. et al. Systematic comparison of constitutive promoters and the doxycycline-inducible promoter. PLoS One 5, e10611 (2010).

47. Pang, Z. et al. In situ identification of cellular drug targets in mammalian tissue. Cell (2022) doi:10.1016/j.cell.2022.03.040.

48. Hao, Y. et al. Integrated analysis of multimodal single-cell data. Cell 184, 3573–3587.e29 (2021).

49. Palla, G., Fischer, D. S., Regev, A. & Theis, F. J. Spatial components of molecular tissue biology. Nat. Biotechnol. 40, 308–318 (2022).

50. McInnes, L., Healy, J. & Melville, J. UMAP: Uniform Manifold Approximation and Projection for Dimension Reduction. arXiv [stat.ML] (2018) doi:10.48550/ARXIV.1802.03426.

## Additional references

51. Wolf, F. A., Angerer, P. & Theis, F. J. SCANPY: large-scale single-cell gene expression data analysis. Genome Biol. 19, 15 (2018).

52. Bradski, G. The openCV library. Dr. Dobb’s Journal: Software Tools for the Professional Programmer 25, 120–123 (2000).

53. Goddard, T. D., Huang, C. C. & Ferrin, T. E. Visualizing density maps with UCSF Chimera. J. Struct. Biol. 157, 281–287 (2007).

54. Hunter, J. D. Matplotlib: A 2D Graphics Environment. Computing in Science & Engineering vol. 9 90–95 (2007).

55. Website. Jones, E., Oliphant, T. & Peterson, P. SciPy: open source scientific tools for Python http://www.scipy.org/ (2001).

56. MacQueen, J. & Others. Some methods for classification and analysis of multivariate observations. in Proceedings of the fifth Berkeley symposium on mathematical statistics and probability vol. 1 281–297 (Oakland, CA, USA, 1967).

57. Higham, D. J. Higham NJ MATLAB guide. Society for Industrial and Applied Mathematics 95–109 (2016).

58. McKinney, W. & Others. Data structures for statistical computing in python. in Proceedings of the 9th Python in Science Conference vol. 445 51–56 (Austin, TX, 2010).

59. Pedregosa, F. et al. Scikit-learn: Machine learning in Python. the Journal of machine Learning research 12, 2825–2830 (2011).

60. Pérez, F., Granger, B. E. & Hunter, J. D. Python: An Ecosystem for Scientific Computing. Computing in Science Engineering 13, 13–21 (2011).

61. Heideman, M., Johnson, D. & Burrus, C. Gauss and the history of the fast fourier transform. IEEE ASSP Magazine 1, 14–21 (1984).

62. van der Walt, S. et al. scikit-image: image processing in Python. PeerJ 2, e453 (2014).

63. Oliphant, T. E. A guide to NumPy. vol. 1 (Trelgol Publishing USA, 2006).

